# COVID-19 lung disease shares driver AT2 cytopathic features with Idiopathic pulmonary fibrosis

**DOI:** 10.1101/2021.11.28.470269

**Authors:** Saptarshi Sinha, Vanessa Castillo, Celia R. Espinoza, Courtney Tindle, Ayden G. Fonseca, Jennifer M. Dan, Gajanan D. Katkar, Soumita Das, Debashis Sahoo, Pradipta Ghosh

**Affiliations:** Department of Cellular and Molecular Medicine, University of California San Diego, La Jolla, CA, 92093; Department of Pediatrics, University of California San Diego, La Jolla, CA, 92093; Department of Computer Science and Engineering, Jacobs School of Engineering, University of California San Diego, La Jolla, CA, 92093; Center for Infectious Disease and Vaccine Research, La Jolla Institute for Immunology (LJI), La Jolla, CA, USA; Department of Medicine, Division of Infectious Diseases and Global Public Health, University of California, San Diego (UCSD), La Jolla, CA, USA; Department of Pathology, University of California San Diego, La Jolla, CA, 92093; Department of Medicine, University of California San Diego, La Jolla, CA, 92093

**Keywords:** Artificial Intelligence/Machine Learning, Immune response, Alveolar Type II Pneumocytes, Damage-associated transient progenitor (DATP), Senescence, ER-stress, Interleukin 15 (IL15)

## Abstract

**Background:** In the aftermath of Covid-19, some patients develop a fibrotic lung disease, i.e., *<u>p</u>*ost-*<u>C</u>*OVID-19 *l*ung *<u>d</u>*isease (PCLD), for which we currently lack insights into pathogenesis, disease models, or treatment options.

**Method:** Using an AI-guided approach, we analyzed > 1000 human lung transcriptomic datasets associated with various lung conditions using two viral pandemic signatures (ViP and sViP) and one covid lung-derived signature. Upon identifying similarities between COVID-19 and idiopathic pulmonary fibrosis (IPF), we subsequently dissected the basis for such similarity from molecular, cytopathic, and immunologic perspectives using a panel of IPF-specific gene signatures, alongside signatures of alveolar type II (AT2) cytopathies and of prognostic monocyte-driven processes that are known drivers of IPF. Transcriptome-derived findings were used to construct protein-protein interaction (PPI) network to identify the major triggers of AT2 dysfunction. Key findings were validated in hamster and human adult lung organoid (ALO) pre-clinical models of COVID-19 using immunohistochemistry and qPCR.

**Findings:** COVID-19 resembles IPF at a fundamental level; it recapitulates the gene expression patterns (ViP and IPF signatures), cytokine storm (IL15-centric), and the AT2 cytopathic changes, e.g., injury, DNA damage, arrest in a transient, damage-induced progenitor state, and senescence-associated secretory phenotype (SASP). These immunocytopathic features were induced in pre-clinical COVID models (ALO and hamster) and reversed with effective anti-CoV-2 therapeutics in hamsters. PPI-network analyses pinpointed ER stress as one of the shared early triggers of both diseases, and IHC studies validated the same in the lungs of deceased subjects with COVID-19 and SARS-CoV-2-challenged hamster lungs. Lungs from *tg***-**mice, in which ER stress is induced specifically in the AT2 cells, faithfully recapitulate the host immune response and alveolar cytopathic changes that are induced by SARS-CoV-2.

**Interpretation:** Like IPF, COVID-19 may be driven by injury-induced ER stress that culminates into progenitor state arrest and SASP in AT2 cells. The ViP signatures in monocytes may be key determinants of prognosis. The insights, signatures, disease models identified here are likely to spur the development of therapies for patients with IPF and other fibrotic interstitial lung diseases.

**Funding:** This work was supported by the National Institutes for Health grants R01-GM138385 and AI155696 and funding from the Tobacco-Related disease Research Program (R01RG3780).

**One Sentence Summary:** Severe COVID-19 triggers cellular processes seen in fibrosing Interstitial Lung Disease

**RESEARCH IN CONTEXT:** *Evidence before this study:* In its aftermath, the COVID-19 pandemic has left many survivors, almost a third of those who recovered, with a mysterious long-haul form of the disease which culminates in a fibrotic form of interstitial lung disease (post-COVID-19 ILD). Post-COVID-19 ILD remains a largely unknown entity. Currently, we lack insights into the core cytopathic features that drive this condition.

*Added value of this study:* Using an AI-guided approach, which involves the use of sets of gene signatures, protein-protein network analysis, and a hamster model of COVID-19, we have revealed here that COVID-19 -lung fibrosis resembles IPF, the most common form of ILD, at a fundamental level—showing similar gene expression patterns in the lungs and blood, and dysfunctional AT2 processes (ER stress, telomere instability, progenitor cell arrest, and senescence). These findings are insightful because AT2 cells are known to contain an elegant quality control network to respond to intrinsic or extrinsic stress; a failure of such quality control results in diverse cellular phenotypes, of which ER stress appears to be a point of convergence, which appears to be sufficient to drive downstream fibrotic remodeling in the lung.

*Implications of all the available evidence:* Because unbiased computational methods identified the shared fundamental aspects of gene expression and cellular processes between COVID-19 and IPF, the impact of our findings is likely to go beyond COVID-19 or any viral pandemic. The insights, tools (disease models, gene signatures, and biomarkers), and mechanisms identified here are likely to spur the development of therapies for patients with IPF and, other fibrotic interstitial lung diseases, all of whom have limited or no treatment options. To dissect the validated prognostic biomarkers to assess and track the risk of pulmonary fibrosis and develop therapeutics to halt fibrogenic progression.

## INTRODUCTION

As the acute phase of the COVID-19 pandemic winds down, the chronic diseases in its aftermath have begun to emerge. For example, many survivors are suffering from a mysterious long-haul form of the disease which culminates in a fibrotic form of interstitial lung disease (post-COVID-19 ILD)(1–9). The actual prevalence of post-COVID-19 ILD (henceforth, *PCLD*) is still emerging; early analysis indicates that more than a third of the survivors develop fibrotic abnormalities. One of the important determinants for PCLD(^10^) is the duration of disease; ∼4% of patients with a disease duration of less than 1 week, ∼24% of patients with a disease duration between 1-3 weeks, and ∼61% of patients with a disease duration > 3 weeks developed fibrosis. Disability among survivors of severe COVID-19 has been primarily attributed to reduced lung capacities (11). While COVID-19– affected lungs and post-Influenza lungs share the histologic pattern of diffuse alveolar damage (DAD)(12) with perivascular T-cell infiltration, key differences have been observed (13). The lungs from patients with Covid-19 also showed (13): (i) widespread thrombosis with microangiopathy; (ii) severe endothelial injury associated with the presence of intracellular virus and disrupted cell membranes; and (iii) neovascularization predominantly through a mechanism of intussusceptive angiogenesis. These findings highlight that despite being a viral pandemic, COVID-19, but not influenza impacts the lungs in ways that go beyond acute DAD, a common phenomenon in progressive ILDs (12). Although a recent study (14) implicated pathogenic subsets of respiratory CD8^+^ T cells in contributing to persistent tissue microenvironment in PCLD, what remains largely unknown are some of the earliest cellular and molecular mechanisms that fuel the progression of fibrosis in COVID-19 lung (but not influenza).

As for how the fibrotic sequel of COVID-19 is managed at present, based on the shared pathological features of overt fibrosis in both end-stage COVID-19 and ILDs, lung transplantation remains the mainstay option for these patients (15–17). Although corticosteroids appear to improve the risk of PCLD(2), beyond that, there is no existing therapeutic option.

Here we seek to unravel the fundamental molecular mechanisms underlying PCLD, identifying key disease drivers (cellular processes, immune pathways, and the signaling cascades that support those pathways). We use artificial intelligence (AI) and machine learning derived gene signatures, the *<u>Vi</u>*ral *<u>P</u>*andemic, ViP and, severe(s) ViP signatures that are induced in all respiratory viral pandemics(18). Besides the ViP signatures, we also use gene signatures induced in the lungs of patients with severe COVID-19. These approaches, in conjunction with experimental validation (in pre-clinical disease models and in human lung tissue), not only helped identify which lung pathology shares fundamental molecular features with COVID-19 lung but also revealed key mechanistic insights into the pathogenesis of PCLD. The findings also provide clues into how to navigate, prognosticate and recapitulate in pre-clinical models this emergent mysterious condition. The objectivity and precision of the AI-guided unbiased approaches enhance the translational potential of our findings.

## METHODS

### KEY RESOURCE TABLE

| **MATERIALS & REAGENTS** |  |  |  |
| --- | --- | --- | --- |
| **ANTIBODIES USED FOR IMMUNOCYTOCHEMISTRY** |  |  |  |
| **Name** | **Manufacturer** | **Catalog number** | **Dilution factor** |
| **p14 ARF (E3X6D)** | Cell Signaling | 74560S | 1:250 |
| **GRP78/BIP Rabbit polyclonal antibody** | Proteintech | 11587-1-AP | 1:500 |
| **P53 Rabbit polyclonal antibody** | Proteintech | 21891-1-AP | 1:200 |
| **Cytokeratin 8 Rabbit polyclonal antibody** | Proteintech | 10384-1-AP | 1:2000 |
| **Claudin 4-specific Rabbit polyclonal antibody** | Proteintech | 16195-1-AP | 1:200 |
| **p21 Waf1 / Cip1 (12D1)** | Cell Signaling | 2947S | 1:50 |
| **CD14** | Invitrogen | MA5-32248 | 1:100 |
| **SARS-CoV2 Nucleoprotein** | Sino Biological | 40143-MM05 | 1:250 |
| **anti-Rabbit IgG Secondary Antibody; AlexaFluor 594 nm** | Invitrogen | A-11012 | 1:500 |
| **anti-Mouse IgG Secondary Antibody, AlexaFluor 488 nm** | Invitrogen | A-31560 | 1:500 |
| **INSTRUMENTS** |  |  |  |
| **Leica DMI4000B (Automated Inverted Microscope)** | Leica Microsystems | DMI4000B |  |
| **Power Pressure Cooker XL** | Tristar Products |  |  |
| **MESO QuickPlex SQ 120** | PBL Assay Science |  |  |
| **SOFTWARE & PACKAGES** |  |  |  |
| **ImageJ** | https://imagej.nih.gov/ij/index.html |  |  |
| **GraphPad Prism** | https://www.graphpad.com/scientific-software/prism/ |  |  |
| **MSD® DISCOVERY WORKBENCH 4.0** | https://www.mesoscale.com/en/products_and_services/software/ |  |  |
| **NetworkX** | https://networkx.org |  |  |
| **Cytoscape** | https://cytoscape.org |  |  |
| **KITS, ENZYMES, CHEMICALS, AND REAGENTS** |  |  |  |
| **DAPI** | Invitrogen | D1306 | 1:500 |
| **Zinc Formalin** | Fisher Scientific | 23-313096 |  |
| **Xylene** | VWR | XX0060-4 |  |
| **Hematoxylin** | BioGenex | HK100-9K |  |
| **Two-Component DAB** | BioGenex | HK542-XAKE | 2 drops per 1mL |
| **Ethanol** | Koptec | UN1170 |  |
| **Sodium Citrate** | Sigma-Aldrich | W302600 |  |
| **Tris Base** | Fisher Scientific | BP154-1 |  |
| **Ethylenediaminetetraacetic acid (EDTA)** | Fischer Scientific | BP118-500 |  |
| **3% Hydrogen Peroxide** | Target | 245-07-3628 |  |
| **ImmPRESS HRP Horse Anti-Rabbit IgG Polymer Detection Kit, Peroxidase** | Vector Labs | MP-7401 |  |
| **Goat serum 2.5%** | Vector Laboratories | S-10120 |  |
| **Normal Antibody Diluent** | MP Biomedicals, LLC | 980641 |  |

### REAGENT VALIDATION

There are no cell lines used in this work. All the antibodies used for immunocytochemistry in this work have been previously validated by commercial entities (in many instances, by confirming loss of signal by target depletion). These details are included in **Supplemental Information 8** (Reagent Validation file).

### DETAILED METHODS

#### Computational Methods

##### StepMiner analysis

*StepMiner* is an algorithm that identifies step-wise transitions using step function in a time-series data(19). *StepMiner* undergoes an adaptive regression scheme to verify the best possible up and down steps based on sum-of-square errors. The steps are placed between time points at the sharpest change between expression levels, which gives us the information about timing of the gene expression-switching event. To fit a step function, the algorithm evaluates all possible steps for each position and computes the average of the values on both sides of a step for the constant segments. An adaptive regression scheme is used that chooses the step positions that minimize the square error with the fitted data. Finally, a regression test statistic is computed as follows:

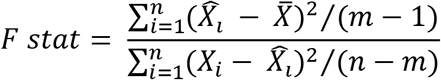

Where *X*_i_ for *i* = 1 to *n* are the values, 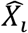 for *i* = 1 to *n* are fitted values. m is the degrees of freedom used for the adaptive regression analysis. 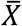 is the average of all the values: 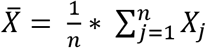. For a step position at k, the fitted values 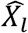 are computed by using 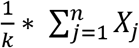 for *i* = 1 to *k* and 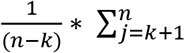.

##### Composite gene signature analysis using Boolean Network Explorer (BoNE)

Boolean network explorer (BoNE) provides an integrated platform for the construction, visualization and querying of a gene expression signature underlying a disease or a biological process in three steps (**Fig 2**, **4, 6D, 8D, S1**): First, the expression levels of all genes in these datasets were converted to binary values (high or low) using the StepMiner algorithm. Second, Gene expression values were normalized according to a modified Z-score approach centered around *StepMiner* threshold (formula = (expr - SThr)/3*stddev). Third, the normalized expression values for every genes were added together to create the final score for the gene signature. The samples were ordered based on the final signature score. Classification of sample categories using this ordering is measured by ROC-AUC (Receiver Operating Characteristics Area Under The Curve) values. Welch’s Two Sample t-test (unpaired, unequal variance (equal_var=False), and unequal sample size) parameters were used to compare the differential signature score in different sample categories. Violin, Swarm and Bubble plots are created using python seaborn package version 0.10.1. Pathway analysis of gene lists were carried out via the reactome database and algorithm(20).

**Figure 1.**
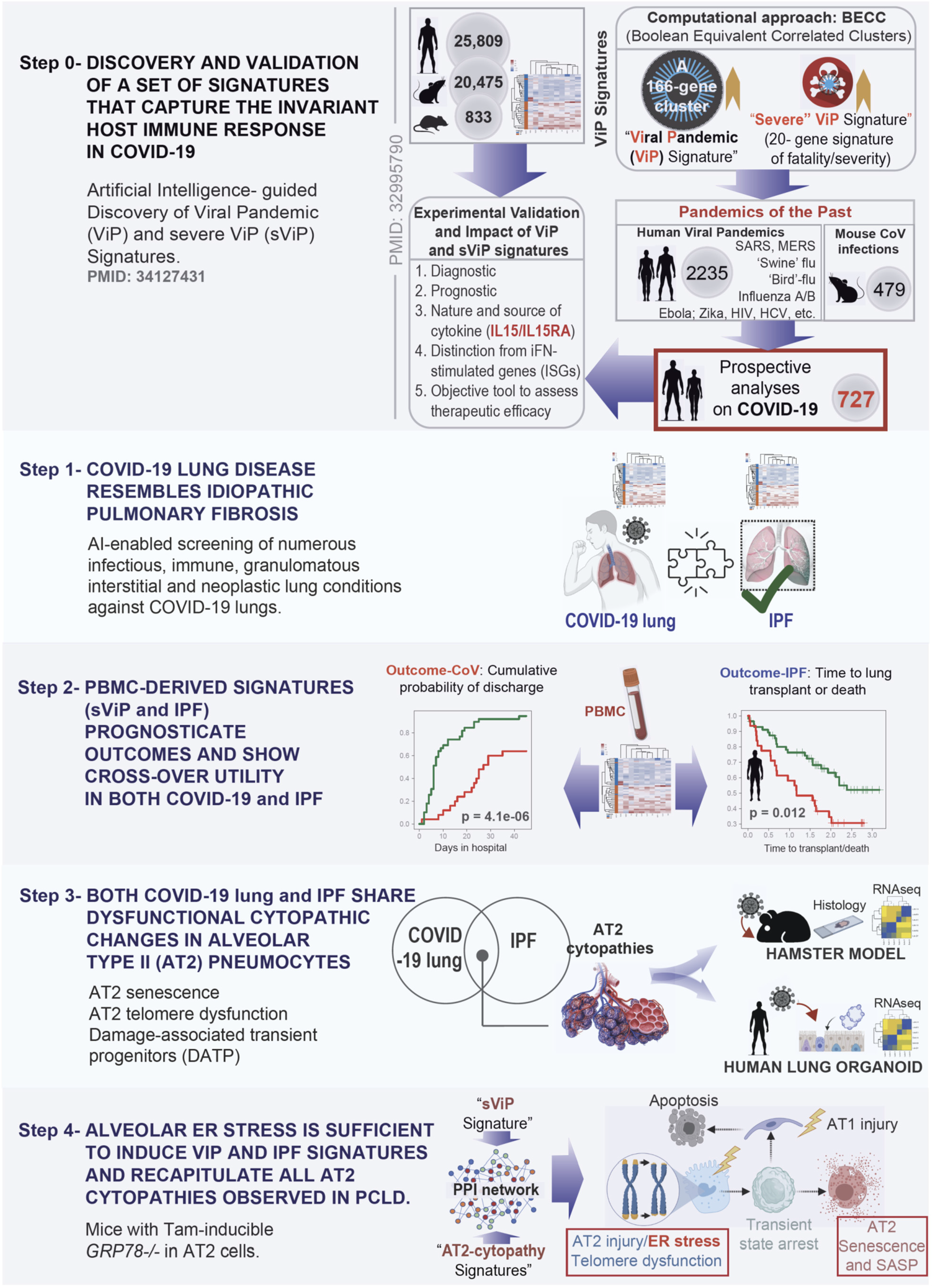
Study design: Artificial Intelligence-guided navigation of COVID-19 lung disease. (*From top to bottom*) *<u>Step 0</u>*: Over 45,000 human, mouse, and rat gene expression databases were mined using machine learning tools called Boolean Equivalent Correlated Clusters (BECC(133)) to identify invariant host response to viral pandemics (ViP). In the absence of a sufficiently large number of COVID-19 datasets at the onset of the COVID-19 pandemic, these ViP signatures were trained on only two datasets from the pandemics of the past (Influenza and avian flu; <u>GSE47963</u>, n = 438; <u>GSE113211</u>, n = 118) and used without further training to prospectively analyze the samples from the current pandemic (i.e., COVID-19; n = 727 samples from diverse datasets). A subset of 20-genes classified disease severity called severe-ViP (sViP) signature. The ViP signatures appeared to capture the ‘invariant’ host response, i.e., the shared fundamental nature of the host immune response induced by all viral pandemics, including COVID-19. *<u>Step1</u>*: The set of ViP/sViP signatures and a CoV-lung specific(13) gene signature was analyzed on diverse transcriptomic datasets representing a plethora of lung diseases; these efforts identified COVID-19 lung disease to be the closest to Idiopathic pulmonary fibrosis (IPF); both conditions induced a common array of gene signatures. *<u>Step 2</u>*: Clinically useful whole-blood and PBMC-derived prognostic signatures previously validated in IPF(27) showed crossover efficacy in COVID-19, and *vice versa*. *<u>Step 3</u>*: Gene signatures of alveolar type II (AT2) cytopathic changes that are known to fuel IPF were analyzed in COVID-19 lung, and predicted shared features were validated in human and hamster lungs and lung-organoid derived models. *<u>Step 4:</u>* Protein-protein interaction (PPI) network built using sViP and AT2 cytopathy-related signatures was analyzed to pinpoint ER stress as a major shared feature in COVID-19 lung disease and IPF, which was subsequently validated in human and hamster lungs.

**Figure 2.**
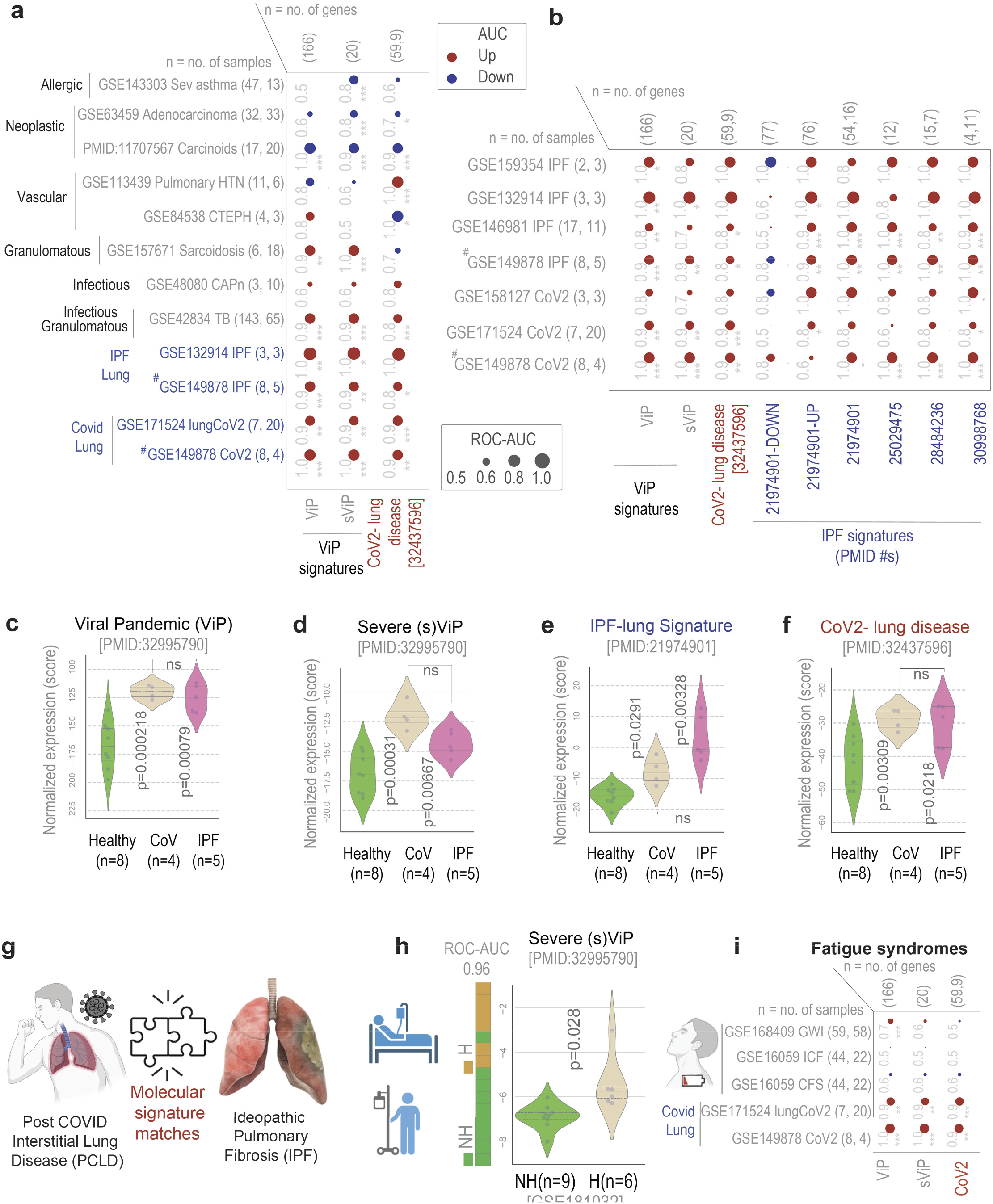
Identification of lung condition(s) that resemble COVID-19 lung disease. a. Bubble plots show the ROC-AUC values (radii of circles are based on the ROC-AUC) show the direction of gene regulation (Up, red; Down, blue) for sample classification based on the induction of ViP/sViP(18) and a previously published human CoV-lung signatures(13) in diverse datasets representing diverse lung pathologies, including COVID-19. Tuberculosis, TB; community acquired pneumonia, CAPn; pulmonary hypertension, PHT; chronic thromboembolic pulmonary hypertension, CTEPH; idiopathic pulmonary fibrosis, IPF; CoV, COVID-19. **b.** Bubble plots in B show the ROC-AUC values for sample classification based on the induction of ViP signatures and several previously published IPF signatures in COVID-19 infected lungs and sc-RNA seq datasets from IPF explants. **c-f.** Violin plots display the extent of induction of ViP (E) or sViP (F), IPF (G) or CoV lung (H) signatures in a pooled cohort of healthy, CoV and IPF lung samples (<u>GSE149878</u>; <u>GSE122960</u>). Welch’s two sample unpaired t-test is performed on the composite gene signature score (z-score of normalized tpm count) to compute the *p* values. In multi-group setting each group is compared to the first control group and only significant *p* values are displayed. **g.** Schematic summarizes the predicted molecular match between COVID-19 lung and IPF based on the crossover validation of gene signatures showcased in B-F. **h.** Bar plots show the ability of sViP signature to classify COVID-19 patients based on hospitalization status and violin plots show the degree of induction of the signature within each group (H, hospitalized; NH, non-hospitalized). Welch’s two sample unpaired t-test is performed on the composite gene signature score (z-score of normalized tpm count) to compute the *p values*. **i.** Bubble plots of ROC-AUC values (radii of circles are based on the ROC-AUC) show the direction of gene regulation (Up, red; Down, blue) for the classification of healthy vs diseased samples based on the ViP/sViP and the CoV-lung signatures on three well known fatigue conditions (blood/PBMC samples from GWI, gulf war illness; CFS, chronic fatigue syndrome). The significance of the AUC was determined by Welch’s t-test; * = p < 0.05, ** = p < 0.01, and *** = p < 0.001. (# denotes a pooled cohort of <u>GSE149878</u>; <u>GSE122960</u>).

**Figure 3.**
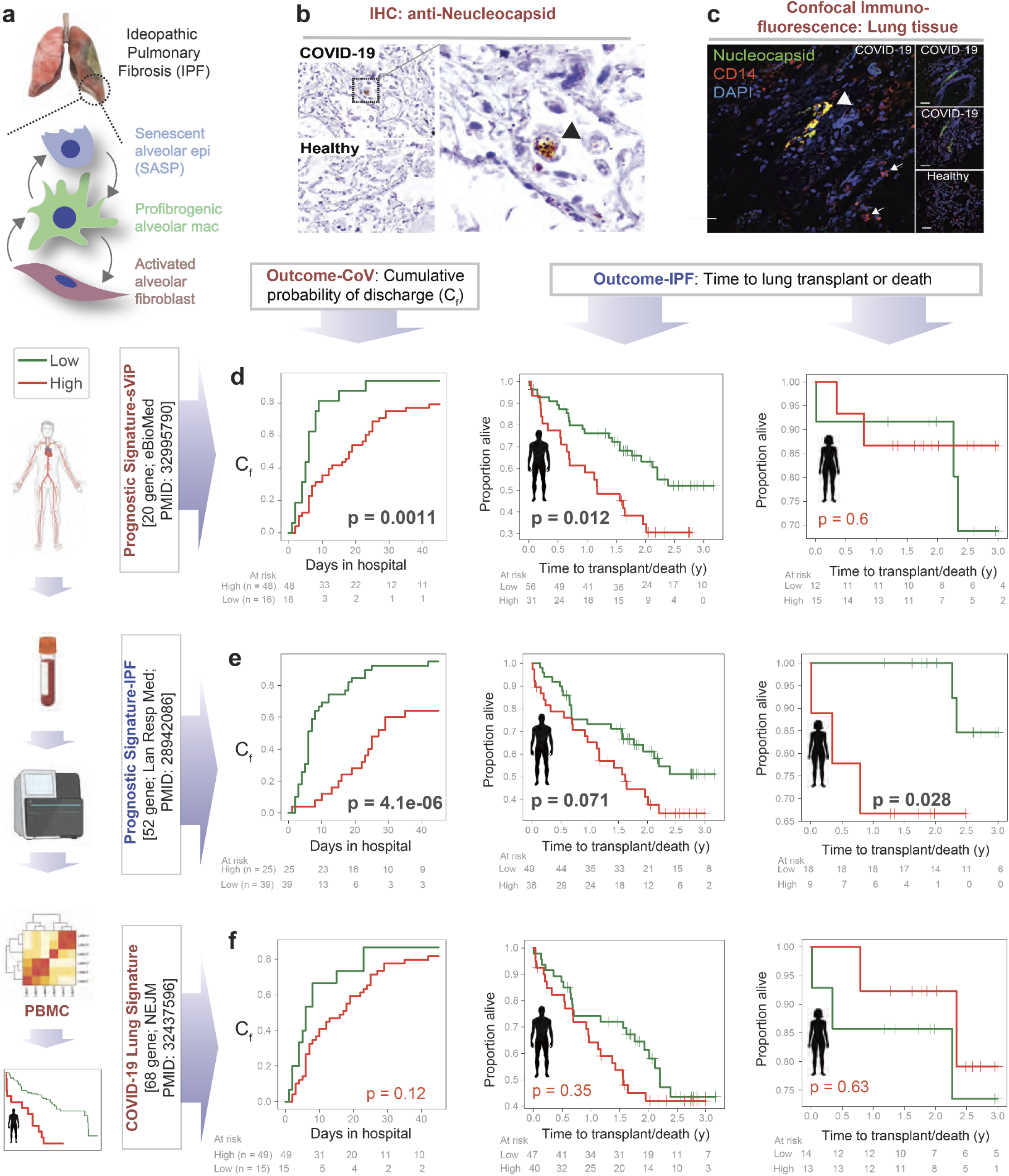
PBMC-derived prognostic signatures in IPF also prognosticate outcome in COVID-19. **a.** Schematic showing the currentunderstanding of the complex interplay between senescent alveolar pneumocyte (AT2), reactive alveolar macrophage, and fibroblasts in the pathogenesis of pulmonary fibrosis. **b-c**. FFPE tissues from normal lungs and from the lungs of subjects deceased due to COVID-19 were analyzed for the SARS-CoV2 viral nucleocapsid protein either by IHC (B) or by confocal immunofluorescence (C). Lungs were co-stained also with CD14 (a marker for activated alveolar macrophage(61)) in C. **d.** Kaplan-Meier plots show the stratification of patients based on the degree of induction of the sViP signature in PBMCs (low-green and high-red groups; determined using the *StepMiner* algorithm(19) and its relationship to either the cumulative probability of discharge of COVID-19 patients (D-*left*; <u>GSE157103</u>; derived from the hospital-free days during a 45-day follow-up), or the time to lung transplant or death in male (D-*middle*) and female (D-*right*) IPF patients (<u>GSE28221</u>). Statistical significances were estimated using the log rank test. **e.** Kaplan-Meier plots on the same cohorts as in D, using a PBMC-derived 52-gene IPF-specific prognostic signature that was previously validated on numerous independent IPF cohorts(27). **f**. Kaplan-Meier plots on the same cohorts as in D-E, using a CoV-lung derived signature.

**Figure 4.**
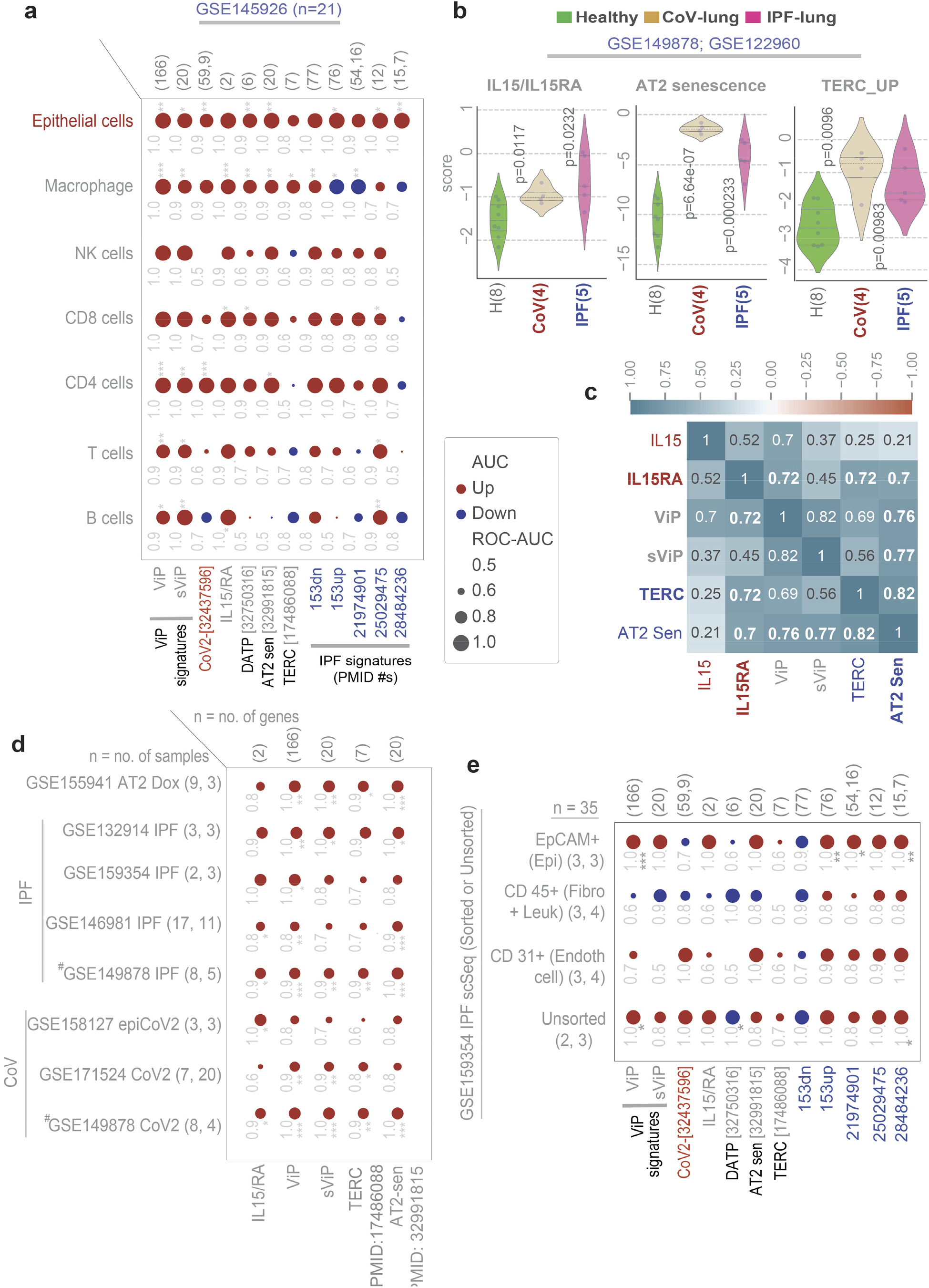
Identification of shared AT2 cytopathies in COVID-19 and IPF. **a**. Bubble plot of ROC-AUC values (radii of circles are based on the ROC-AUC) demonstrating the direction of gene regulation (Up, red; Down, blue) for the classification of various cell types between healthy and CoV lung based on various ViP/sViP and IPF gene signatures in **Fig 2a-b** alongside *IL15/IL15RA* and additional signatures of AT2 cytopathies that are encountered and implicated in IPF (DATP, damage associated transient progenitors; Sens, senescence; TERC, *<u>Te</u>*lomerase *<u>R</u>*NA *<u>C</u>*omponent). Numbers in parenthesis indicate PMIDs. **b.** Violin plots display the extent of induction of a *IL15/IL15RA* composite score (B-*left*), or a signature of AT2 senescence(70) (B-*middle*), or a gene set for telomere dysfunction (JU_AGING_TERC(134)) in a pooled cohort of healthy, CoV and IPF lung samples (<u>GSE149878</u>; <u>GSE122960</u>). Welch’s two sample unpaired t-test is performed on the composite gene signature score (z-score of normalized tpm count) to compute the *p* values. In multi-group setting each group is compared to the first control group and only significant p values are displayed. **C.** Correlation matrix showing the Pearson’s correlation coefficient between different gene signatures in the pooled healthy, CoV and IPF lung dataset (<u>GSE149878</u>; <u>GSE122960</u>). **d.** Bubble plots of ROC-AUC values (radii of circles are based on the ROC-AUC) demonstrating the direction of gene regulation (Up, red; Down, blue) for the classification based on AT2 senescence-related signatures on other publicly available CoV and IPF bulk RNA seq datasets. **e.** Bubble plots of ROC-AUC values (radii of circles are based on the ROC-AUC) demonstrating the direction of gene regulation (Up, red; Down, blue) for different types of cells in an independent sc-RNA seq datasets of IPF explants and healthy lung tissues. The significance of the AUC was determined by Welch’s t-test; * = p < 0.05, ** = p < 0.01, and *** = p < 0.001. (# denotes a pooled cohort of <u>GSE149878</u>; <u>GSE122960</u>).

**Figure 5.**
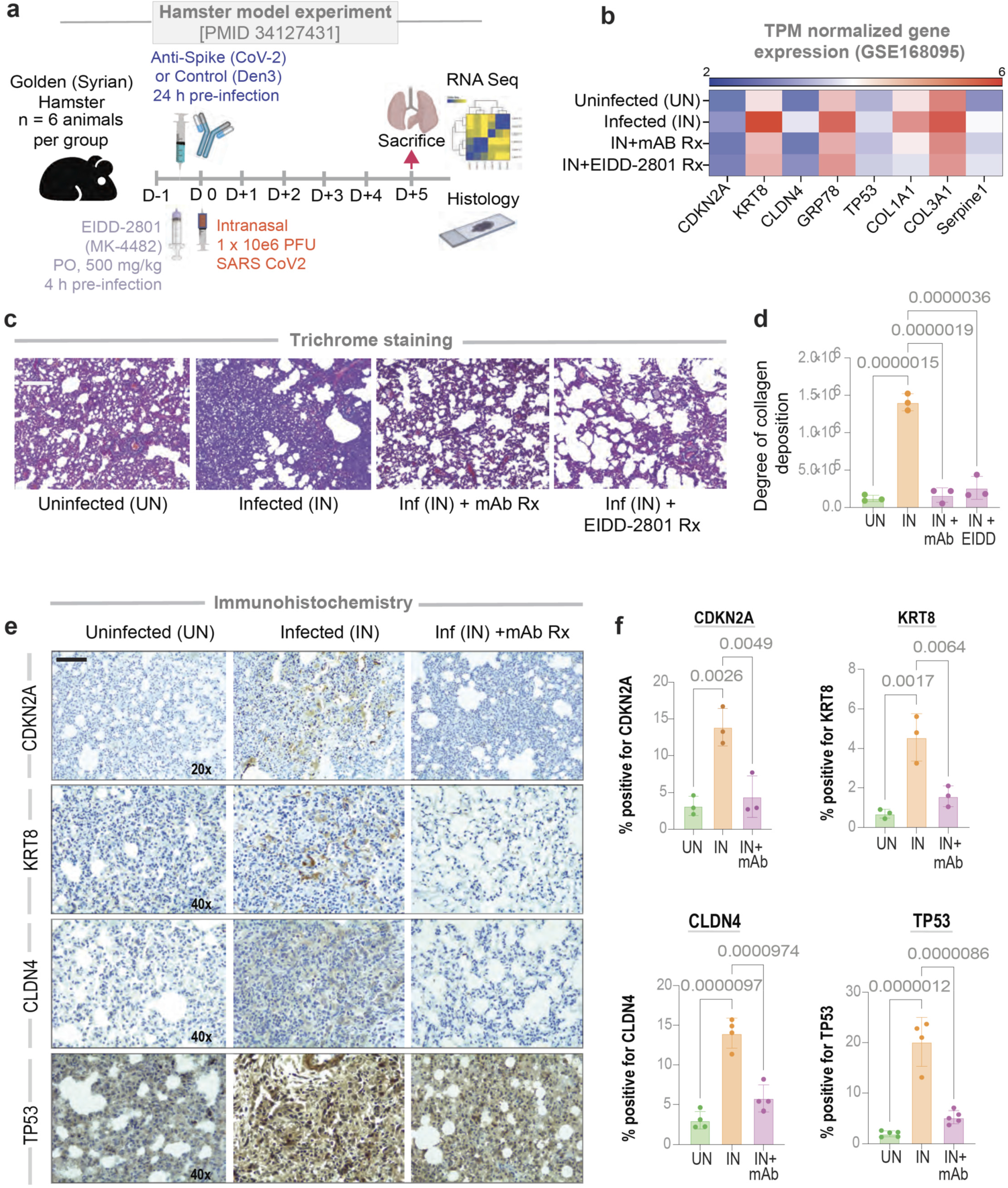
SARS-CoV-2-challenged hamster lungs faithfully recapitulate AT2 cytopathies in COVID-19. **a.** Schematic showing the experimental design for COVID-19 modeling in golden Syrian hamsters. *Uninf*, uninfected; *Den3 and Anti-CoV-2* indicate SARS-CoV-2 challenged groups that received either a control mAb or the clone CC12.2 of anti-CoV-2 IgG(31), respectively. PFU, plaque-forming units. **b**. Heatmap showing transcripts/million (TPM)-normalized gene expression of specific genes associated with AT2 senescence (*CDKN2A*, *TP53*), damage associated transition state progenitor (DATP) arrested state (*CLDN4*, *KRT8*), ER stress (*HSPA5*/*GRP78*) and SASP-state (*SERPINE1*, *COL1A*, *COL3A*). **c-d**. Lungs harvested from the uninfected, infected, and infected but anti-CoV-2 IgG and infected but EIDD-2801 treated hamsters have been analyzed for the abundance of collagen deposition by Trichrome staining and quantified by ImageJ. Representative images are shown panel c. Scale bar = 200 µm. Results of the quantification are displayed as bar plot in D. Statistical significance was analyzed by one way ANOVA. Error bars represent S.E.M; n = 3. **e-f**. Lungs harvested from the uninfected, infected, and infected but anti-CoV-2 IgG groups were analyzed by IHC for the indicated proteins and quantified by ImageJ-IHC profiler. Representative images are shown panel E. Scale bar = 200 µm. Results of the quantification are displayed as violin plots in F. Statistical significance was analyzed by one way ANOVA; n = 4.

**Figure 6.**
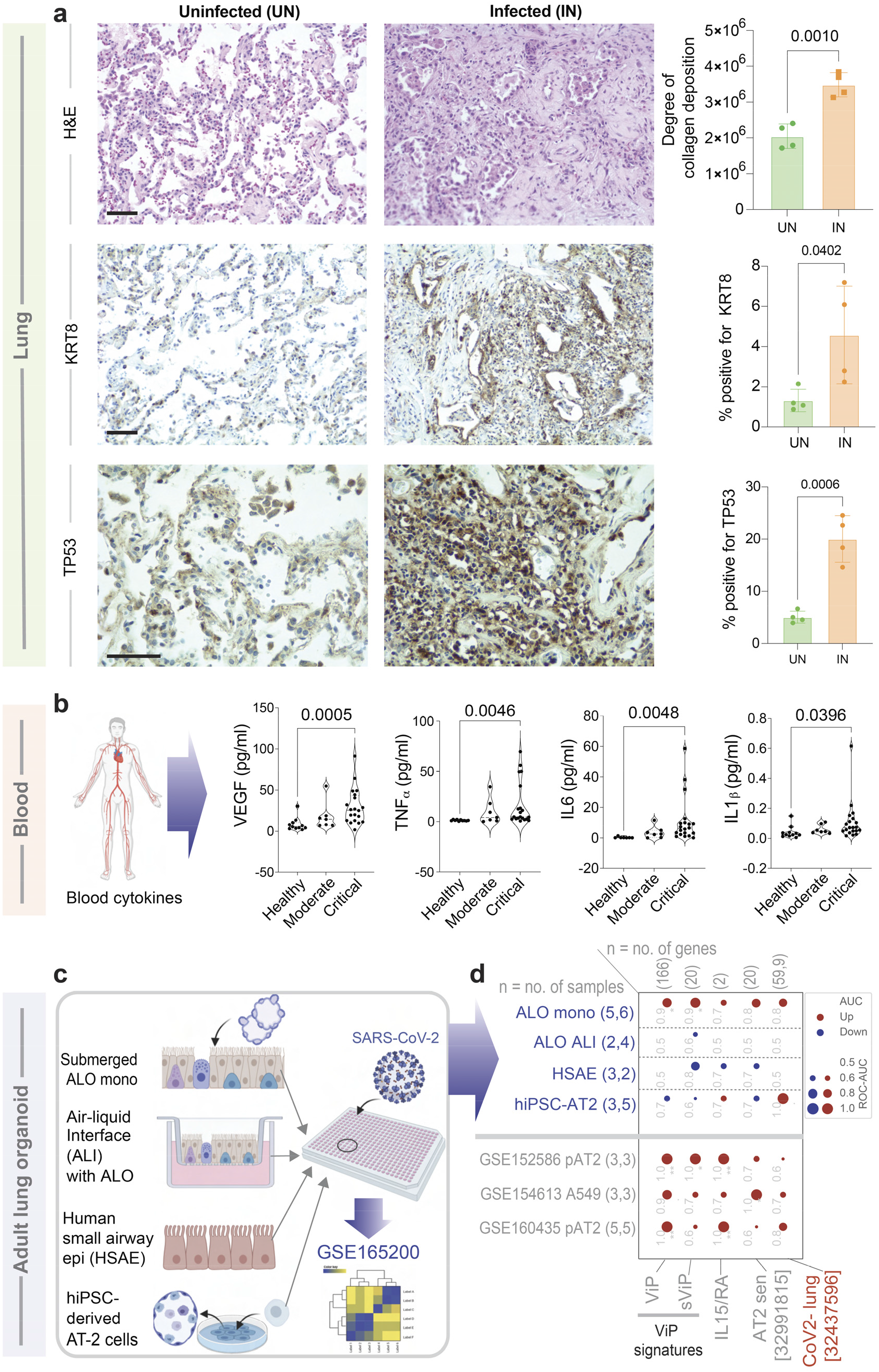
AT2 cytopathic features of IPF are detected in COVID-19 lung tissues and human pre-clinical organoid models of COVID-19. **a**. FFPE lung tissues collected during rapid autopsy of deceased COVID-19-infected subjects and histologically normal surgically resected lung tissues were analyzed by H&E staining (*top*) and for the expression of KRT8 (a marker of DATP(67); *middle*) and TP53 (a marker of senescence; *bottom*) by IHC. Representative fields are shown on the left. Scale bar = 200 µm. Images were quantified by IHC profiler (ImageJ) and displayed as violin plots on the right. **b**. Violin plots display the abundance of key senescence-associated cytokines in the plasma from adults with COVID-19 and healthy volunteers. Statistical significance was analyzed by unpaired t-test. See also **Supplemental Information 3-4** for patient demographics and cytokine concentrations. **c-d**. Schematic in C summarizes the various pre-clinical *in vitro* models of the human lung that were challenged with SARS-CoV-2, followed by RNA sequencing(28) (*top* rows; blue font; D). Other independent SARS-CoV-2-challenged alveolar pneumocyte datasets are listed in grey font (D). Bubble plots in D show the ROC-AUC values (radii of circles are based on the ROC-AUC) demonstrating the direction of gene regulation (Up, red; Down, blue) for the classification of uninfected vs infected samples based on the signatures (below).

**Figure 7.**
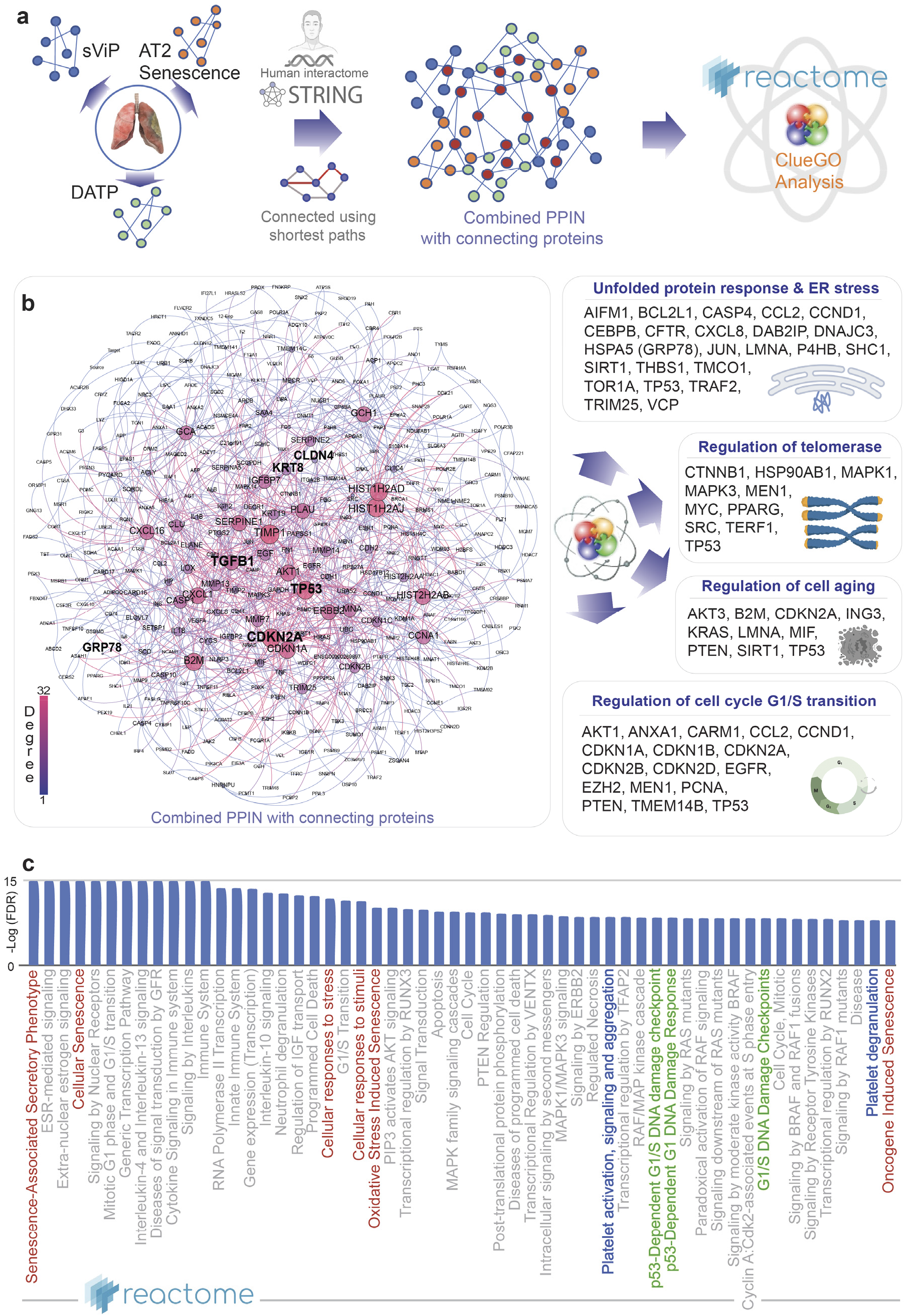
Protein-protein interaction network (PPIN) analysis predicts ER stress in AT2 cells as a driver pathophysiology in COVID-19. **a**. Schematic summarizes the key steps in building a PPI network using as ‘seeds’ the genes in the sVIP, AT2 senescence and transition state arrest (DATP) signatures and connecting them using shortest paths prior to carrying out network analyses. **b**. The combined PPI network is shown on the left and the degree of connectivity is indicated. See also **Supplemental Information 5** for the complete list of nodes sorted by their degrees of connectivity. Schematics on the right summarize the results of CluGo analysis, which identifies cellular processes and key genes that may perpetuate lung fibrosis. **c**. Reactome pathway analysis of the PPI network lists the most significant pathways that are associated with the node list in the network in B. Red = senescence related processes; Green = DNA damage response pathways; Blue = Platelet activation and degranulation.

**Figure 8.**
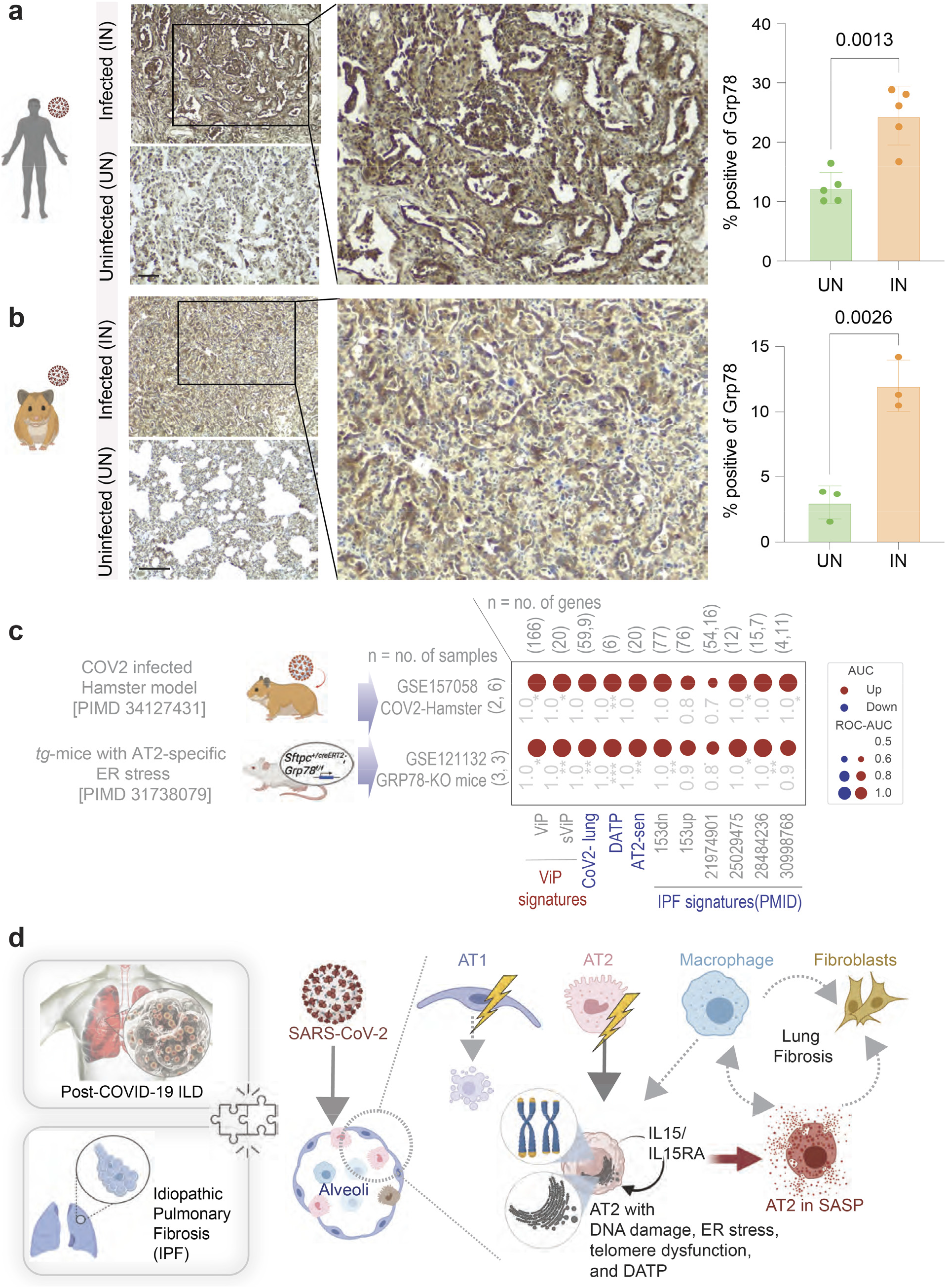
Induction of ER stress in AT2 cells is observed in COVID-19 and is sufficient to mimic the host immune response in COVID-19 and IPF. **a-b**. FFPE uninfected (control) and SARS-CoV-2-infected human (A) and hamster (B) lungs were analyzed for GRP78 expression by IHC. Representative fields are shown on the left. Scale bar = 200 µm. Images were quantified by IHC profiler (ImageJ) and displayed as violin plots on the right. **c**. Bubble plots show the ROC-AUC values (radii of circles are based on the ROC-AUC) demonstrating the direction of gene regulation (Up, red; Down, blue) for the classification of WT vs *GRP78*-KO murine lung (bottom row) and uninfected vs. infected hamster lung (top row) samples based on the signatures (below). **d**. Schematic summarizes the findings in this work and the proposed working model for the contributions of various alveolar cells in fueling the fibrotic progression in both IPF and COVID-19. The significance of the AUC was determined by Welch’s t-test; * = p < 0.05, ** = p < 0.01, and *** = p < 0.001.

##### ViP and severe (s)ViP signatures

ViP (Viral Pandemic) signature(18) is derived from a list of 166 genes using Boolean Analysis of large viral infection datasets (training datasets: <u>GSE47963,</u> n = 438; <u>GSE113211,</u> n = 118). This 166-gene signature was conserved in all viral pandemics, including COVID-19, inspiring the nomenclatures ViP signature. A subset of 20-genes classified disease severity called severe-ViP signature using an additional cohort (<u>GSE101702</u>, n = 159)(18). No interstitial lung disease (ILD) datasets, or for that matter any other lung dataset was used during the process of discovery and validation of the ViP/sViP signatures. To compute the ViP signature, first the genes present in this list were normalized according to a modified Z-score approach centered around *StepMiner* threshold (formula = (expr -SThr)/3*stddev). The normalized expression values for every probeset for 166 genes were added together to create the final ViP signature. The severe ViP signature is computed similarly using 20 genes. The samples were ordered finally based on both the ViP and severe-ViP signature. A color-coded bar plot is combined with a violin plot to visualize the gene signature-based classification.

##### Protein-protein interaction network (PPIN) construction

PPIN has been constructed using the gene signatures as a set of seed nodes. The nodes between the seed nodes were fetched using the connecting shortest paths and their components from the human protein interaction dataset of the STRING database(21). A high cutoff of STRING interaction score has been chosen based on the proteins present in the signature list to neglect the false positive interactions.

##### Transcriptomic datasets and data analysis

Several publicly available microarrays and RNASeq databases were downloaded from the National Center for Biotechnology Information (NCBI) Gene Expression Omnibus (GEO) server (22–24). Gene expression summarization was performed by normalizing Affymetrix platforms by RMA (Robust Multichip Average) and RNASeq platforms by computing TPM (Transcripts Per Millions) (25, 26) values whenever normalized data were not available in GEO. We used log2(TPM +1) as the final gene expression value for analyses. GEO accession numbers are reported in figures and text. A catalog of all datasets analyzed in this work can be found in **Supplemental Information 1**. Patient demographics, where available, are curated in **Supplemental Information 6** and show that age, gender, and ethnicity are reflective of a wide population of patients impacted by the COVID-19 pandemic hence, the findings and conclusions of the current study may maintain relevance in a similar population. *StepMiner* analysis and methodologies for composite gene signatures, outcome analyses, and violin plots are detailed in Supplementary Online Methods.

##### Survival outcome analysis

Kaplan-Meier analysis was done for three different gene signatures: sViP (18), 52 gene IPF signature (27) and COVID-lung signature (13) over two different survival datasets of COVID patients (<u>GSE157103</u>) and IPF patients (<u>GSE28221</u>). Hospital-free days analysis (45 days followup) of COVID-19 patients (<u>GSE157103</u>) were limited to less than 70 years old patients. The high and low group are separated based on *StepMiner* threshold on the composite score of the gene expression values. The IPF dataset (<u>GSE28221</u>) included 114 patients with outcome was divided into male (n = 87) and female (n = 27) population for checking the gender bias. The survival data for the IPF dataset was also censored at 3 years for comparative analysis. The statistical significance of KM plots was assessed by log rank test.

##### Single Cell RNA Seq analysis

Single Cell RNASeq data from <u>GSE145926</u> was downloaded from Gene Expression Omnibus (GEO) in the HDF5 Feature Barcode Matrix Format. The filtered barcode data matrix was processed using Seurat v3 R package. B cells (CD19, MS4A1, CD79A), T cells (CD3D, CD3E, CD3G), CD4 T cells (CCR7, CD4, IL7R, FOXP3, IL2RA), CD8 T cells (CD8A, CD8B), Natural killer cells (KLRF1), Macrophages, Monocytes and DCs (TYROBP, FCER1G), Epithelial (SFTPA1, SFTPB, AGER, AQP4, SFTPC, SCGB3A2, KRT5, CYP2F1, CCDC153, TPPP3) cells were identified using relevant gene markers using SCINA algorithm. Pseudo bulk datasets were prepared by adding counts from the different cell subtypes and normalized using log2(CPM+1). All other single-cell RNA Seq datasets <u>GSE159354</u>, <u>GSE132914</u>, <u>GSE146981,</u> and <u>GSE149878</u> were processed using scanpy python package (v1.5.1) and converted to pseudo-bulk samples by adding counts from all cells and normalized using log2(CPM+1). The downstream BoNE composite score analyses were performed similarly to regular bulk RNASeq dataset.

##### Statistical analyses

Gene signature is used to classify sample categories and the performance of the multi-class classification is measured by ROC-AUC (Receiver Operating Characteristics Area Under the Curve) values. A color-coded bar plot is combined with a density plot to visualize the gene signature-based classification.All statistical tests were performed using R version 3.2.3 (2015-12-10). Standard t-tests were performed using python scipy.stats.ttest_ind package (version 0.19.0) with Welch’s Two Sample t-test (unpaired, unequalvariance (equal_var=False), and unequal sample size) parameters. Multiple hypothesis correction was performed by adjusting *p* values with statsmodels.stats.multitest.multipletests (fdr_bh: Benjamini/Hochberg principles). Sample number of each analysis is provided with associated plots beside each GSE ID no. or sample name. Pathway analysis of gene lists were carried out via the Reactome database and CluGo algorithm. Reactome identifies signaling and metabolic molecules and organizes their relations into biological pathways and processes. Kaplan-Meier analysis is performed using lifelines python package version 0.14.6. The statistical significance of KM plots was assessed by log-rank test. Violin, Swarm and Bubble plots are created using python seaborn package version 0.10.1.

##### Multivariate analysis

Multivariate regression has been performed on the single cell (<u>GSE132914</u>) and bulk (<u>GSE150910</u>) sequence IPF datasets of various covid and IPF signatures (in Figure S1). Multivariate analysis models the healthy vs IPF samples as a linear combination of composite scores of sViP, ViP, CoV-lung and six independent IPF signatures. Here, the <u>statsmodels</u> module from python has been used to perform Ordinary least-squares (OLS) regression analysis of each of the variables. The choice of these datasets was driven by the criteria that they are high quality datasets with maximal unique patient samples. For single cell dataset (<u>GSE132914</u>), cell types has been considered as an additional variable. The p-value for each term tests the null hypothesis that the coefficient is equal to zero (no effect).

#### Experimental Methods

##### Rapid autopsy procedure for tissue collection

The lung specimens from the COVID-19 positive human subjects were collected as described in detail previously (18, 28) using autopsy (study was IRB Exempt). All donations to this trial were obtained after telephone consent followed by written email confirmation with next of kin/power of attorney per California state law (no in-person visitation could be allowed into our COVID-19 ICU during the pandemic). The team member followed the CDC guidelines for COVID-19 and the autopsy procedures^8, 9^. Lung specimens were collected in 10% Zinc-formalin and stored for 72 h before processing for histology. Autopsy #2 was a standard autopsy performed by anatomical pathology in the BSL3 autopsy suite. The patient expired and his family consented for autopsy. After 48 hours, lungs were removed and immersion fixed whole in 10% formalin for 48 hours and then processed further. Lungs were only partially fixed at this time (about 50% fixed in thicker segments) and were sectioned further into small 2-4cm chunks and immersed in 10% formalin for further investigation. Autopsies #4 and #5 were collected from rapid postmortem lung biopsies. The procedure was performed in the Jacobs Medical Center ICU (all the ICU rooms have a pressure-negative environment, with air exhausted through HEPA filters [Biosafety Level 3 (BSL3)] for isolation of SARS-CoV-2 virus). Biopsies were performed 2-4 hours after patient expiration. Ventilator was shut off to reduce aerosolization of viral particles at least 1 hour after loss of pulse and before the sample collection. Every team member had personal protective equipment in accordance with the University policies for procedures on patients with COVID-19 (N95 mask + surgical mask, hairnet, full face shield, surgical gowns, double surgical gloves, booties). Lung biopsies were obtained after L-thoracotomy in the 5th intercostal space by our cardiothoracic surgery team. Samples were taken from the left upper lobe (LUL) and left lower lobe (LLL) and then sectioned further.

The histologically confirmed normal lung samples (non-COVID-19 control) were obtained from surgically removed lobes for indications such as neoplastic and non-neoplastic lung nodules, at sites distant from the pathologic lesion.

##### Immunofluorescence of normal and COVID-19 lung tissue samples

FFP-embedded lung tissue sections underwent antigen retrieval immersed in Tris-EDTA buffer (pH 9.0) and boiled at 100°C inside a pressure cooker for 10 min. Once sections returned to room temperature, samples were washed in deionized (DI) water and then permeabilized and blocked for 2 h using a blocking buffer (2 mg/mL BSA and 0.1% Triton X-100 in PBS); as described previously(29). Primary antibodies CD14 (1:100 dilution) and SARS-COV2 Nucleoprotein (1:250 dilution) were diluted in blocking buffer and allowed to incubate overnight at 4°C. Secondary antibodies and DAPI (1:500 dilution) were diluted in blocking buffer and allowed to incubate for 2 h in the dark. ProLong Glass was used as a mounting medium. #1 Thick Coverslips were applied to slides and sealed. Samples were stored at 4°C until imaged. As negative control (to determine the specificity of staining) we eliminated the step of incubation with primary antibodies while maintaining all other steps.

Images were acquired at room temperature with Leica TCS SPE confocal and with DMI4000 B microscope using the Leica LAS-AF Software. Images were taken with a 40× oil-immersion objective using 405-, 488-, 561-nm laser lines for excitation. Z-stack images were acquired by successive Z-slices of 1µm in the desired confocal channels. Fields of view that were of interest and demonstrated a positive immunofluorescence signal were selected and imaged. Z-slices of a Z-stack were overlaid to create maximum intensity projection images; all images were processed using FIJI (Image J) software. Power analyses for these studies have been presented in **Supplemental Information 7**.

##### Collection of blood from COVID-19 patients

Blood from COVID-19 donors was either obtained at a UC San Diego Health clinic under the approved IRB protocols of the University of California, San Diego (UCSD; 200236X) or recruited at the La Jolla Institute under IRB approved (LJI; VD-214). COVID-19 donors were California residents, who were either referred to the study by a health care provider or self-referred. Blood was collected in acid citrate dextrose (ACD) tubes (UCSD) or in EDTA tubes (LJI) and stored at room temperature prior to processing for plasma collection. Seropositivity against SARS-CoV-2 was confirmed by ELISA. At the time of enrollment, all COVID-19 donors provided written informed consent to participate in the present and future studies. Patient characteristic is listed in **Supplemental Information 3**.

##### Human serum cytokines measurement

Human serum cytokines measurement was performed using the V-PLEX Custom Human Biomarkers from MSD platform [MESO QuickPlex SQ 120]. Human serum samples fractionated from peripheral blood of KD and MIS-C patients (all samples collected prior to the initiation of treatments) were analyzed using customized standard multiplex plates as per the manufacturer’s instructions using recommended software [MSD® DISCOVERY WORKBENCH 4.0].

##### COVID-19 lung models in-a-dish

Monolayers derived from adult lung organoids (ALOs), primary airway cells, or hiPSC-derived alveolar type II (AT2) pneumocytes were infected with SARS-CoV-2 to create in vitro lung models of COVID-19 as described previously(28). The raw data and processed data were deposited in Gene Expression Omnibus under accession no. <u>GSE157057</u>. These datasets were analyzed here using various gene signatures.

##### COVID-19 modeling in Syrian hamsters

Lung samples from 8-week-old Syrian hamsters were generated from experiments conducted exactly as in a previously published study (18, 30, 31). Briefly, the infected (IN) arm was challenged with an intranasal dose of 10^6^ PFU SARS-CoV-2 on day 1. The indicated treatments (or corresponding vehicle controls) were administrated either a day prior (monoclonal antibodies; 2 mg per animal, intraperitoneal route) or 4 h prior (EIDD-2801; 500 mg/kg, oral route) to the CoV-2 challenge as a single dose. Animals were scarified 5^th^ day after infection to collect blood and lung tissues (18, 30, 31). Viral titers, weight and lung histology findings have been published elsewhere (18, 30, 31). Animal studies were approved and performed in accordance with Scripps Research IACUC Protocol #20-0003 (30, 31). Power analyses for these studies have been presented in **Supplemental Information 7**.

##### Immunohistochemistry

COVID-19 samples were inactivated by storing in 10% formalin for 2 days and then transferred to zinc-formalin solution for another 3 days. The deactivated tissues were transferred to 70% ethanol and cassettes were prepared for tissue sectioning. The slides containing hamster and human lung tissue sections were deparaffinized in xylene (Sigma-Aldrich Inc., MO, USA; catalog# 534056) and rehydrated in graded alcohols to water. For GRP78/ BIP, p53, Cytokeratin 8, and Claudin 4-specific polyclonal antigen retrieval, slides were immersed in Tris-EDTA buffer (pH 9.0) and boiled for 10 minutes at 100°C. Slides were immersed in Sodium Citrate Buffer (pH 6.0) and boiled for 10 minutes at 100°C, for p14 ARF (E3X6D) and p21 Waf1/Cip1(12D1) antigen retrieval. Endogenous peroxidase activity was blocked by incubation with 3% H2O2 for 5-10 minutes. To block non-specific protein binding either 2.5% goat or 2.5% horse serum (Vector Laboratories, Burlingame, USA; catalog# MP-7401 or S-1012) was added. Tissues were then incubated with the following antibodies: rabbit GRP78/BIP polyclonal antibody (1:500 dilution; proteintech®, Rosemont, IL, USA; catalog# 11587-1-AP), rabbit p53 polyclonal antibody (1:200 dilution ; proteintech®, Rosemont, IL, USA; catalog# 21891-1-AP), rabbit Cytokeratin 8 polyclonal antibody (1:2000 dilution; proteintech®, Rosemont, IL, USA; catalog# 10384-1-AP), and rabbit Claudin 4-specific polyclonal antibody (1:200 dilution, proteintech®, Rosemont, IL, USA; catalog# 16195-1-AP) for 1.5 hours at room temperature in a humidified chamber then rinsed with TBS twice for 3 minutes each. Antibodies were prepared in antibody diluent (MP Biomedicals, LLC., Catalog # 980641) for rabbit p14 ARF (E3X6D) monoclonal antibody (1:250 dilution; Cell Signaling Technology, Danvers, MA, USA; catalog# 74560S) and rabbit p21 Waf1/Cip1 (12D1) monoclonal antibody (1:50 dilution; Cell Signaling Technology, Danvers, MA, USA; catalog# 2947S) that were then incubated in a humidified chamber overnight at 4°C, then rinsed with TBST twice for 5 minutes each. Sections were incubated with horse anti-rabbit (Vector Laboratories, Burlingame, USA; catalog# MP-7401) secondary antibodies for 30 minutes at room temperature in a humidified chamber and then washed with TBS or TBST 3x, 5 minutes each. Then incubated with DAB (BioGenex, Fremont, CA, USA; catalog # HK542-XAKE) for 5-10 minutes and counterstained with hematoxylin (BioGenex, Fremont, CA, USA; catalog# HK100-9K), dehydrated in graded alcohols, cleared in xylene, and cover slipped. All sides were visualized by Leica DM1000 LED (Leica Microsystems, Germany).

##### IHC quantification

IHC images were randomly sampled at different 300×300 pixel regions of interest (ROI). The ROIs were analyzed using IHC Profiler(32). IHC Profiler uses a spectral deconvolution method of DAB/hematoxylin color spectra by using optimized optical density vectors of the color deconvolution plugin for proper separation of the DAB color spectra. The histogram of the DAB intensity was divided into 4 zones: high positive (0 to 60), positive (61 to 120), low positive (121 to 180) and negative (181 to 235). High positive, positive, and low positive percentages were combined to compute the final percentage positive for each region of interest (ROI). The range of values for the percent positive is compared among different experimental groups. To calculate the statistical significance (determination of the p value), one way ANOVA has been performed between respective groups of samples.

##### Quantification of collagen

Due to acidophilic nature of collagen, it stains with eosin in H&E stanning. The relative extracellular collagen deposition was assessed using color deconvolution in ImageJ. The same color deconvolution technique was also used when collagen was stained using Masson’s Trichrome (33, 34). To calculate the statistical significance (determination of the p value), one way ANOVA has been performed between respective groups of samples.

##### Ethics statement

Animal studies were performed in compliance with the ethical guidelines outlined in a Scripps Research Institutional Animal Care and Use Committee (IACUC)-approved protocol (Protocol #20–0003; PI: Tom Rogers) (31).

Blood from COVID-19 donors was either obtained at a UC San Diego Health clinic under the approved IRB protocols of the University of California, San Diego (UCSD; 200236X) or recruited at the La Jolla Institute under IRB approved (LJI; VD-214). An informed consent was obtained in all cases. COVID-19 donors were California residents who were either referred to the study by a health care provider or self-referred.

The lung specimens from the COVID 19 positive human subjects were collected using autopsy (study was IRB Exempt). All donations to this trial were obtained after telephone consent followed by written email confirmation with next of kin/power of attorney per California state law (no in-person visitation could be allowed into our COVID-19 ICU during the pandemic). The team member followed the CDC guidelines for COVID19 and the autopsy procedures.

## RESULTS

### A study design that uses gene signatures as a computational framework to navigate COVID-19 lung disease

Because PCLD is still an emergent illness that lacks insights into disease pathophysiology, we resorted to a study design (**Figure 1**) that is geared to achieve three goals: (i) maximize rigor by using diverse transcriptomic datasets cohorts (a total of 1825 unique samples, human: 1766; mouse: 41, hamster: 18; see **Supplemental Information 1**), but (ii) minimize bias by using well characterized sets of gene signatures from independent publications (see **Supplemental Information 2**), and (iii) retain objectivity and precision by using artificial intelligence/machine learning (AI/ML) tools that are supported by precise mathematical algorithms and statistical tools (see *Methods*).

First, to ensure that the findings maintain relevance to respiratory viral pandemics and COVID-19, we used a recently discovered 166-gene signature that was found to be conserved in all *<u>vi</u>*ral *<u>p</u>*andemics (ViP), including COVID-19, and a subset of 20-genes within that signature that classifies disease severity (18); the latter was found to be consistently and significantly induced in the most severe cases of COVID-19 and prognostic of poor outcome, e.g., death or invasive respiratory support (see **Supplemental Information 2**). These ViP signatures were rigorously validated on ∼45000 datasets representing the pandemics of the past (Training datasets: Influenza and avian flu; <u>GSE47963</u>, n = 438; <u>GSE113211</u>, n = 118; Validated in numerous datasets as cited before) and used without further training to prospectively analyze the samples from the current pandemic (i.e., COVID-19) (**Figure 1**, *Step 0*). The ViP signatures appeared to capture the ‘invariant’ host response, i.e., the shared fundamental nature of the host immune response induced by all viral pandemics, including COVID-19. Since they were discovered, these signatures have helped navigate the emergent syndrome of multisystem inflammatory syndrome in children (MIS-C; a.k.a. PIMS-TS)(35). To avoid overreliance on one set of signatures (i.e., ViP/sViP, which are tissue-agnostic signatures of host response), and increase the specificity for lung tissue, an independent gene signature was included, one that is induced in COVID-19 lung tissues, but not in influenza (13).

We began first by using the ViP signatures as a starting computational framework to navigate the syndrome of COVID-19 lung disease, in conjunction with another independently reported (13) COVID-19-lung derived signature to identify idiopathic pulmonary fibrosis (IPF)(36) as a computational match for COVID-19 lung disease (**Figure 1**, *Step 1*). The subsequent steps in the study use the ViP signatures alongside IPF-specific PBMC-derived prognostic signatures (**Figure 1**, *Step 2*) and signatures of alveolar cytopathic changes (**Figure 1**, *Step 3*) to systematically assess the degree of similarities between COVID-19 lung and IPF and the crossover of pathophysiologic processes in the two conditions. In the final step (**Figure 1**, *Step 4*), a set of gene signature-inspired protein-protein interaction (PPI) network analysis is used which pinpointed alveolar ER stress as a potential early step in both IPF and PCLD, which is sufficient to recapitulate not just the ViP signatures, but also the alveolar cytopathic features that are shared between the diseases. Key predictions are validated in human lung tissues and plasma from COVID-19 subjects, hamster models of COVID-19, and human pre-clinical lung models (**Figure 1**, *Steps 3-4*).

### COVID-19 lung disease and IPF induce a common set of gene expression signatures

We first asked which pathologic lung condition comes closest to COVID-19 lung disease regarding the host immune response. To this end, we used the ViP/sViP signatures (found to be induced in all CoV samples tested so far (18)) and a second CoV-lung disease gene signature which was derived from a differential expression analysis on lung samples from healthy controls vs. fatal COVID-19(13) (see **Supplemental Information 2**). As for the lung diseases, we ensured that all four major pathologic conditions were represented: neoplastic (e.g., carcinoids and adenocarcinoma), granulomatous (e.g., tuberculosis/TB and sarcoidosis), allergic/infectious (e.g., severe asthma and community-acquired pneumonia/CAP and tuberculosis) and vasculopathic (e.g., pulmonary hypertension/PHT and chronic thromboembolic pulmonary hypertension/CTEPH) (**Figure 2a**). As expected, ViP/sViP signatures were induced in infectious diseases (CAP and tuberculosis), and to our surprise, these signatures were upregulated also in sarcoidosis, a granulomatous condition that progresses to fibrosis in ∼20% of the patients (37, 38) (**Figure 2a**). Only two diseases (tuberculosis and IPF) showed significant induction of both ViP and CoV-lung signatures, of which only one condition (IPF) manifests as a diffuse disease with patchy parenchymal involvement resembling COVID-19 lung disease (**Figure 2a**).

We next analyzed a set of six signatures, each independently derived from diverse IPF-centric studies (39–42), on diverse CoV samples; we also analyzed the ViP/sViP(18) and CoV-lung (13) signatures on 4 independent single-cell RNA sequence (scRNA seq) datasets from IPF explants. We found that CoV samples induced the IPF signatures and IPF samples induced ViP/CoV-lung signatures (**Figure 2b**). It is noteworthy that such induction of ViP/sViP signatures was undetectable in the bulk RNA seq studies on IPF explants (**Supplementary Figure 1a**); this is in keeping with numerous reports that scRNA-seq obtains cell-level resolution in IPF that bulk-RNA seq misses(43–45).

Next, we created a multivariate model to decompose the covariance between ViP, sViP, CoV lung and all other IPF signatures to estimate the amount of overlaps between their induction in the IPF samples. These studies showed that the ViP signature (alongside a some IPF signatures) is an independent variable that is induced in IPF samples (**Supplementary Figure 1b**). However, such induction was observed exclusively in scSeq datasets (**Supplementary Figure 1b**), but not in bulk Seq dataset (**Supplementary Figure 1c**), which is also consistent with **Supplementary Figure 1a**.

Finally, in a pooled dataset (<u>GSE149878</u>)(46) that included both CoV and IPF samples the ViP (**Figure 2c**), sViP (**Figure 2d**), IPF-lung (**Figure 2e**), and CoV-lung (**Figure 2f**) were indistinguishably induced in both samples compared to healthy controls. These results indicate that CoV and IPF signatures have crossover utility for sample classification and indicate that IPF is the closest computational match to COVID-19 lung disease (**Figure 2g**). Consistent with the fact that PCLD occurs in survivors of severe COVID-19 that required hospitalization(47), we found that sViP signature was induced significantly higher in hospitalized compared to non-hospitalized subjects (**Figure 2h**). Finally, we asked what, if any relationship exists between another poorly understood feature of long-haul COVID-19, i.e., severe fatigue(48, 49). To improve rigor, we analyzed a set of other fatigue syndromes (e.g., gulf war illness, chronic fatigue syndrome, and idiopathic chronic fatigue) and for consistency, restricted the analysis to only whole blood/PBMC samples (**Figure 2i**). ViP/sViP and CoV-lung immune signatures were induced only in CoV samples, but not in the remaining fatigue-associated syndromes (**Figure 2i**), indicating that gene expression signatures in long-haul COVID-19 are distinct from other fatigue-associated syndromes. Instead, fatigue in long-haul COVID-19 may be of similar reasons/pathophysiology that has been observed in IPF (50). It is noteworthy that profound fatigue is a shared phenomenon in both IPF and sarcoidosis (50, 51), both carry the risk of progressive fibrosis, and both conditions induce the ViP/sViP response (**Figure 2a**). These results show that COVID-19 lung is closest to IPF: both are characterized by lung fibrosis, and share gene expression patterns, suggestive of shared molecular and cellular mechanisms of disease pathogenesis.

### PBMC-derived prognostic signatures in COVID-19 and IPF show crossover utility in both diseases

Previously we showed using sc-RNA seq as well as bulk-RNA seq on sorted cells, that ViP/sViP signatures are primarily induced in COVID-19 by lung epithelial cells and myeloid cells (both alveolar macrophages and PBMCs)(18). When assessed in whole blood and/or PBMCs, the sViP signature was prognostic in COVID-19— a high expression of this 20-gene signature was associated with prolonged hospital stay and mechanical ventilation (18). We noted that a PBMC-based 52-gene signature has previously been validated in numerous IPF datasets (27). In the setting of IPF, the role of PBMC-derived lung macrophages in the activation of alveolar fibroblasts has been cemented by several studies (52–57) (**Figure 3a**); the activated macrophages and myofibroblasts are thought to cross-stimulate each other, resulting in a vicious cycle that assures propagation of fibrosis throughout the lung in end-stage IPF (52, 58). Emerging studies on CoV lung have also reported the presence of pro-fibrogenic alveolar macrophages in severe COVID-19 (15, 59, 60). Because monocytes/macrophages induce ViP signatures, whereas fibroblasts do not (18, 30, 31), we focused on monocytes and investigated if the PBMC-derived prognostic signatures for IPF also prognosticate outcome in CoV, and *vice versa*.

We found that alveolar macrophages (marked by CD-14 (61)) in the lungs of deceased subjects contained the viral nucleocapsid protein (**Figure 3b-c**); this is consistent with observations reported by others (25, 62). The 20-gene sViP signature was prognostic in the blood datasets from both COVID-19 (**Figure 3d**; *left*) and IPF subjects (**Figure 3d**; *middle, right*); among IPF subjects, the signature was prognostic exclusively in male subjects (**Figure 3d**; *middle*). The 52-gene ‘IPF-specific’ prognostic signature (27) retained its ability to prognosticate outcomes in both male and female patients with IPF (as expected; **Figure 3e**; *middle, right*), but also did so in the setting of COVID-19 (**Figure 3e**; *left*). By contrast, the CoV-lung tissue-derived signature that is known to distinguish CoV from both healthy and influenza-affected lungs(63), did not prognosticate outcome in either COVID-19 (**Figure 3f**; *left*) or IPF subjects (**Figure 3f**; *middle vs. right*). It indicates that the PBMC-derived signatures in CoV and IPF retain their prognostic relevance in both conditions, with at least one key notable difference; sViP, but not the IPF signature revealed the well-established gender specific differences in the progression of IPF. These results indicate that PBMCs in both IPF and COVID-19 have similar gene induction patterns and that the cross-over prognostic value of IPF- and sViP signatures induced in the PBMCs may reflect the profibrogenic immune response that is shared in both conditions.

### Alveolar type II (AT2) telomere dysfunction and senescence are shared features in both COVID-19 and IPF

Besides the contributions from PBMCs, a contemporary paradigm in IPF is that chronic injury to distal lung tissue leads to either loss or altered function of epithelial stem cells (i.e., AT2 cells), which promote dysregulated repair and pathogenic activation of fibroblasts (64–66). We asked if the AT2 dysfunction states previously reported in IPF also occur in COVID-19. To this end, we analyzed a single cell dataset of bronchoalveolar lavage from COVID-19 patients (<u>GSE145926</u>) for the induction of ViP/sViP, CoV-lung, and IPF signatures and other signatures that are indicative of three important AT2 cytopathies: (i) Damage associated transient progenitor (DATP(67)), a distinct *KRT8*+/*CLDN4*+ AT2 lineage that has been independently shown, nearly simultaneously by 3 independent groups(67–69) to be associated with activated myofibroblasts during lung fibrosis in the setting of IPF(66). (ii) AT2-senescence signature, which was derived from another IPF-focused study (70) that used human and mouse models to pinpoint the Doxycycline-induced program of p53-dependent cellular senescence, AT2 cell depletion, and spontaneous, progressive pulmonary fibrosis; and (iii) Aging-associated telomerase dysfunction (TERC_UP) signature, which was derived from aging telomerase knockout (*Terc*^-/-^) mice. We found that the entire panel of signatures analyzed were upregulated, most consistently, only in the epithelial cells (**Figure 4a**), whereas only some were induced and not others in the non-epithelial compartments.

We next confirmed that a composite score of *IL15/IL15RA*, AT2-senescence, and telomerase dysfunction (TERC_UP) signatures were induced indistinguishably in both CoV and IPF samples compared to healthy controls (**Figure 4b**). A correlation matrix (**Figure 4c**) on the entire cohort showed that while both IL15 and IL15RA correlate with the ViP signature (as expected), IL15RA, but not IL15 correlate with TERC_UP and AT2-senescence signatures. Both ViP and sViP signatures correlate with AT2 senescence. Furthermore, when we tested AT2-senescence and telomere dysfunction in other CoV datasets, we found that these features were conserved in most CoV samples across diverse cohorts (**Figure 4d**). An independent sc-RNA seq IPF dataset further confirmed that, much like CoV lung, the induction of ViP/sViP in the alveolar epithelium (EpCAM+; **Figure 4e**) is associated with IL15/IL15RA induction, AT2-senescence, and telomere dysfunction. These findings support the existence of telomere dysfunction and DNA damage in both acute COVID-19 and IPF.

### Hamsters, but not mice recapitulate the host immune response and AT2 cytopathic features of COVID-19

Evidence suggests that there is a wide gap between COVID-19 in humans and animal models(71); interspecies-related differences, such as host specificity and divergent immune responses can lead to misinterpretation. To objectively assess which model improves the translational potential of any discovery from this study, we curated all publicly available transcriptomic datasets from pre-clinical rodent models of COVID-19 and analyzed them for the induction of the ViP/sViP, IL15/IL15RA composite score, CoV-lung, IPF signatures and signatures of AT2 senescence and transition-state arrest (DATP) (**Supplementary Figure S2a**). We saw that the golden Syrian hamsters, but not the humanized transgenic mouse models (**Supplementary Figure S2a**; *bottom*) induce most, if not all the key signatures upon SARS-CoV-2 challenge (**Supplementary Figure S2a**; *top*). Most importantly, all signatures (**Supplementary Figure S2a**; *top*), including that of AT2-damage associated transient progenitor population (**Supplementary Figure S2b**; DATP) were effectively reversed with two therapeutic approaches. The first approach was the use of N-hydroxycytidine, the parent of the prodrug MK-4482 (Molnupiravir, EIDD-2801) which has not only proven as a potent and selective oral antiviral nucleoside analogue in mice, guinea pigs, ferrets and human airway epithelium organoids (71–76) but also showed promise in Phase IIa trials in the treatment of COVID-19 patients (NCT04405570(77)). The second approach was the use of SARS-CoV-2-neutralizing antibodies whose design was inspired by monoclonal antibodies (mAbs) isolated from convalescent donors(31). A specific isotype of this antibody, which binds to the receptor-binding domain (RBD-A) of SARS-CoV-2 spike protein in a fashion that precludes binding to host ACE2, was demonstrated as effective in preventing infection and weight loss symptoms, in cell-based and *in vivo* hamster models of infection, respectively.

These findings show that hamster lungs best recapitulate the host immune response, the nature of the cytokine storm (IL15-centric), and the AT2 cytopathies that are shared between COVID-19 and IPF. It is also in keeping with emerging data that the Syrian hamsters best emulate COVID-19(71); they also recapitulate progressive fibrosis in IPF(78).

### Validation of the predicted AT2 cytopathic features of COVID-19 in hamster and human lungs

We next sought to determine if the SARS-CoV-2 virus can induce the AT2 cytopathic changes and whether effective therapeutics can abrogate those changes. We analyzed by RNA seq and trichrome stain the lungs from SARS-CoV-2-challenged golden Syrian hamsters who were pre-treated either with EIDD-2801 or anti-Spike (CoV-2) mAb or untreated controls (see *Methods* and study protocol in **Figure 5a**) (30, 31). Weight loss, histologic confirmation for the presence/extent of inflammatory infiltrates in the lung and serum viral titers in these experimental cohorts have been confirmed and reported previously(18, 30, 31). Key genes associated with AT2 senescence (*Tp53* and *Cdkn2a*), ER stress (*Hspa5/Grp78*), DATP (*Krt8* and *Cldn4*) and fibrosis (*Col1a, Col3a,* and *Serpine1*) were all induced in the infected lungs (compared to uninfected controls) but not in those that were treated (**Figure 5b**). Trichrome staining confirmed that compared to uninfected controls, collagen deposition was induced in the infected lungs, but not in those that were treated (**Figure 5c-d**). Immunohistology studies confirmed that Cdkn2a, Krt8, Cldn4 and Tp53 proteins were expressed in the injured alveoli, but not in the treated lungs (**Figure 5e-f**). Because the degree of viremia, the severity of the infection, and host immune response (the IL15-centric ViP signatures) were significantly blunted by both treatments(18, 30, 31), the orally administered anti-viral EIDD-2801 and the systemic administration of anti-spike mAb, we conclude that the AT2 cytopathic features are a consequence of severe CoV infection.

We next confirmed the extensive deposition of eosin-positive collagenous materials in lung tissues obtained at rapid autopsies on deceased subjects with COVID-19 compared to normal lung tissues (**Figure 6a**; *top*). KRT8 and TP53 were also induced in the COVID-19 lungs (**Figure 6a**; *middle* and *bottom*). Finally, in plasma ELISA studies in a cohort of symptomatic COVID-19 patients who presented to the UC San Diego Medical Center with varying disease severity, ranging from mild to fatal (see **Supplemental Information 3-4**), we detected several cellular senescence-associated cytokines during the acute compared to the convalescent visit (**Figure 6b; Supplementary Figure S3**). It is noteworthy that VEGF, TNFα, IL6, and IL1β, all cytokines that have been implicated in the fibrotic process post-COVID-19 (reviewed in (79)) were significantly elevated only in the most critical form of the disease. Findings are also consistent with the fact that VEGF-targeted therapeutics (Bevacizumab) in COVD-19 patients improved lung oxygenation up to day 28 during follow-up(80).

We next asked if the key aspects of the host immune response and AT2 cytopathic changes are induced in the various human pre-clinical *in vitro* models of COVID-19. We assessed our published models of COVID-19-in-a-dish (28) (**Figure 6c, 6d-***top 4 rows*) for AT2-senescence and for their ability to mimic the unique milieu in COVID-19 lungs(13). We also analyzed CoV2-challenged primary(p)AT2 and A549 cells (**Figure 6d-***bottom 3 rows*). We found that submerged monolayers of adult lung organoids (ALO mono), which were previously found to outperform fetal lung organoids (81) and other models in recapitulating the gene expression signatures observed in human COVID-19 lungs(28), also outperformed here. While most models induced some signatures, the submerged ALO model induced them all (**Figure 6d**).

### Protein-protein network analysis predicts AT2 ER stress as an early initiating event in COVID-19

To identify some of the earliest AT2 cell processes that might be triggered as a consequence of the host immune response in acute COVID-19, which may precede the development of transient state arrest and senescence, we resorted to a PPI network analysis. We used the genes in the sViP (18), AT2 senescence(70) and DATP(67) signatures as ‘seed’ nodes to fetch additional connecting nodes from human protein-protein interactions in STRING database (21) (**Figure 7a;** see **Supplemental Information 5)**. Once built, we assessed the network for degrees of connectivity (**Figure 7b**; *left*) and representation of cellular processes in the most connected nodes by using two approaches: CluGo analysis(82) (**Figure 7b**; *right*) and Reactome pathway analysis(20) (**Figure 7c**). While CluGO analysis showed an overrepresentation of proteins associated with unfolded protein response, ER stress, regulation of telomerase, cell aging and, cell cycle (G1/S phase transition) (**Figure 7b**; *right*), Reactome analysis showed enrichment of pathways associated with senescence (**Figure 7c**; *red*), platelet activation and degranulation (**Figure 7c**; *blue*), and, Tp53-dependent DNA damage response (**Figure 7b**; *green*). That platelet activation (indicative of a prothrombotic state) is captured within the PPI network is in keeping with what appears to be a hallmark of PCLD(13). Numerous independent observations in diverse cohorts of patients have now confirmed that platelet dysfunction amplifies endotheliopathy in COVID-19 (83) and that a vicious cycle of thromboinflammation, endothelial dysfunction, and immunothrombosis are key pathogenic mechanisms in COVID-19(84, 85). It is noteworthy, that we found that levels of IL6 and platelets showed a significant negative correlation (**Supplementary Figure 3**). Because IL6 has been specifically implicated in platelet hyper-reactivity and thrombosis(86), as well as in thrombus resolution (87), the strong negative correlation may suggest the presence of thromboinflammation; the latter is one of the first and most distinctive pathological features of CoV-lung to be reported since the beginning of this pandemic(13).

As for AT2-centric early events/processes, we decided to focus on ER stress because the PPI network revealed so, and because there is fundamental support for its role in cytokine elaboration, apoptosis, and disrupted progenitor cell function as upstream drivers of lung fibrosis(88). AT2 ER stress has been shown to serve as a causal role for profibrogenic epithelial states (SASP) that is shared between aging-associated pulmonary fibrosis and IPF(89), raising the possibility that it may also be a common underlying factor in fibrotic COVID-19 lung disease.

### ER stress in AT2 cells is observed in COVID-19 and is sufficient to recapitulate the host immune response in COVID-19 and IPF

We next sought to investigate if alveolar ER stress is a feature in CoV-lung. To this end, we analyzed infected human (**Figure 8a**) and hamster (**Figure 8b**) lungs for the ER chaperone *Grp78*, a key regulator of ER homeostasis expression by IHC. We found that compared to normal uninfected lungs, infected lungs showed a significant increase in Grp78 staining and that the intensity of such staining was highest in the epithelial cells lining the alveolar spaces in both species (**Figure 8a-b**).

To determine if ER stress in AT2 cells may contribute to the epithelial dysfunction and fibrosis that is observed as a sequel of COVID-19, we leveraged a previously characterized *tg*-mouse model of AT2-specific conditional [tamoxifen (Tmx)-inducible] deletion of Grp78 (*Sftpc^+/creERT2^;Grp78^f/f^*)(89). These mice are known to lose weight and die (more pronounced in male and old mice) due to lung inflammation, and spatially heterogeneous fibrosis characterized by fibroblastic foci and hyperplastic AT2 cells, all features of IPF(89). We found that compared to their littermate controls, lungs from these mice (unlike other transgenic mouse models of COVID-19; see **Figure S2a**) induced the ViP/sViP, CoV2-lung, and IPF signatures, as well as the AT2 cytopathic changes such as DATP and AT2 senescence (**Figure 8c**; *bottom row*). In fact, the pattern of gene expression was similar to SARS-CoV-2-challenged hamster lungs (**Figure 8c**; *top row*). These findings indicate that ER stress is sufficient to faithfully recapitulate the host immune response and alveolar cytopathic changes that are observed in COVID-19 lung disease and IPF.

## DISCUSSION

The major discovery we report here is that lung disease in severe COVID-19 resembles IPF, the most common form of ILD, at a fundamental level—showing similar gene expression patterns (ViP and IPF signatures), shared prognostic signatures (in circulating PBMCs), dysfunctional cell states (AT2 and monocytes), and dysfunctional AT2 processes (ER stress, telomere instability, and senescence)]. It is also noteworthy that this semblance between COVID-19 and IPF was identified through a comprehensive and unbiased computational approach (see **Figure 1**) which compared gene expression signatures in COVID-19 against transcriptomic datasets representing a plethora of neoplastic, granulomatous, allergic/infectious, and vasculopathic pathologic conditions of the lung. Although two other diseases (sarcoidosis and tuberculosis) that are characterized by focal fibrosis or lower incidence of fibrosis were found to induce some of the COVID-19-associated lung signatures, IPF induced them all and matched COVID-19 closely due to the diffuse nature of fibrosis. This finding suggests that the nature of the host immune response to injury and the proximal AT2 cytopathies we report here are unlikely to be mere sequelae of cellular and molecular events during any fibrotic remodeling of the lung and may instead be specific to ILDs such as IPF and COVID-19.

Our findings support the following model (see **Figure 8d**): Upon injury (viral infection/inflammatory cytokines), diffuse alveolar damage is associated with extensive loss of AT1 cells, which are duly replaced by AT2 cells that serve as progenitors that are capable of self-renewal and differentiation into AT1 cells (90–92). But such physiologic regeneration and healing process is impaired when AT2 progenitors are also injured and begin to elicit ER stress responses, which induces telomerase activity (93, 94). Because severe COVID-19 has been linked to short telomere lengths(94) [not as a cause, i.e, SARS-CoV-2 infection does not shorten telomeres, but as a predisposition (95)], it is possible that patients who have short and/or dysfunctional telomeres fail to adequately respond to ER stress, and instead, become senescent (through p53 activation), accumulate DNA damage and enter a dysfunctional *KRT8*+ transitional stem cell state (96). The latter is a phenomenon that has been independently reported by three groups, under different names: PATS, pre-alveolar type-1 transitional cell state (69); ADI, alveolar differentiation intermediates (68); DATP, damage-associated transient progenitors (67). All three groups reported their presence in fibrotic regions of IPF lungs; an increase of AT2 cells in the transitional state was invariably accompanied by an increase in myofibroblasts. Our PPI network analyses suggested that AT2 senescence and progenitor arrested state in the setting of a ViP/sViP-immune response (cytokine storm contributed by PBMCs) may support a SASP phenotype, which is a pathological feature of aging and IPF lung (97, 98).

It is possible that fibroblasts/myofibroblasts also contribute to this vicious cycle of inflammation and AT2 dysfunction. These findings show that these two distinct clinical syndromes, IPF, which predates the current pandemic by many decades, and the novel COVID-19, share a similar profile of host immune response, both in the lung microenvironment (in AT2 cells, to be specific) as well as in the circulating blood/PBMCs. We not only formally define the nature of that host response and provide computational tools to measure the extent of such response, but also chart the cascade of cytopathic changes in the alveoli that are critical for the profibrogenic state. It is noteworthy that the same host immune response is seen also in MIS-C, a new disease that co-emerged with COVID-19, and in KD (which shares overlapping features with MIS-C in clinical presentation (35)).

There are three major impacts and/or implications of the findings reported in this study.

First, our finding that COVID-19 and IPF share fundamental host immune response and alveolar cytopathic features is in keeping with the fact that both diseases share epidemiologic similarities—they primarily impact older adults, males more than females, and are characterized by progressive worsening of dyspnea and lung function (99, 100). The induction of ViP/sViP signatures in IPF is consistent with gathering consensus in the past decade that IPF may be a multi-trigger infection-driven chronic inflammatory condition (36, 101–103). Finally, patients with existing ILDs have greater odds of death due to COVID-19 compared with adults without ILD, even after controlling for age, sex, and comorbidities(104–106). On day 30 of COVID-19, 35% of patients with fibrotic idiopathic ILD had died compared with 19% of those with other ILDs(107). Whether the shared molecular or pathophysiologic features we report here synergize to increase fatality remains unknown; however, these shared features support the rationalization of clinical trials in COVID-19 using FDA-approved drugs for IPF, nintedanib (Ofev®), and pirfenidone (Esbriet®) (<u>NCT04856111</u> and <u>NCT04653831</u>), and anecdotal case reports and case series have already chronicled the benefits of their use in COVID-19 lung disease (108–110).

Second, most severe COVID-19 patients develop pneumonia and hyperinflammation likely related to a macrophage activation syndrome (111) commonly named “cytokine storm”. Although this storm has been implicated, and it is comprised of the generic mix of all the typical cytokines(10), which exact component is the pro-fibrogenic driver remains unclear. By showing that an IL15/IL15RA-centric cytokine storm is the key shared phenomenon between CoV and IPF lungs, this study provides a link between hyperinflammation and the sequelae of fibrosis. Findings are consistent with prior reports of circulating IL15 as a biomarker for prognostication in IPF and other ILDs(112–114). As for what may be contributing to such a cytokine response, myeloid cells are likely to be major culprits, but dysfunctional AT2 cells cannot be ruled out. Prior studies using bleomycin-challenged transgenic mice that lack the *Telomeric Repeat Binding Factor 2* in the AT2 cells (*Trf2^Fl/Fl^;Sftpc-CreER*) have shown that AT2 cells with telomere dysfunction upregulate an IL15-centric pathways(115). Mutations in the two main components of the telomerase holoenzyme complex that is responsible for creating new telomeric DNA, *Telomerase reverse transcriptase (TERT)* or *telomerase RNA (TERC),* are major monogenic causes of pulmonary fibrosis(116). Patients with IPF have shortened telomeres; short telomeres and TERT/TERC-disease associated variants were associated with specific clinical and biological features and reduced transplant-free survival (116, 117). Similarly, short telomeres increase the risk of severe COVID-19(95, 118, 119), pulmonary fibrosis, and poorer outcomes(119, 120). It is possible that injury, DNA damage and IL15 signaling in AT2 cells are one component in the profibrogenic cascade and suggest that IL15-targeted therapeutics may be beneficial in the most severe cases of COVID-19 to prevent fibrotic sequelae.

Third, our finding that shared alveolar cytopathic changes (e.g., DNA damage, progenitor state arrest, SASP, and ER stress) fuel fibrogenic programs in both COVID-19 and IPF are insightful because AT2 cells are known to contain an elegant quality control feedback loop to respond to intrinsic or extrinsic stress; a failure of such quality control results in diverse cellular phenotypes (reviewed in (88)): ER stress (121), defective autophagy, mitochondrial dysfunction, apoptosis, inflammatory cell recruitment, profibrotic signaling, and altered progenitor function (96, 122, 123) that ultimately converge to drive downstream fibrotic remodeling in the lung. Prior work (70), which led to the discovery of the AT2-senescence signature (which we used here), had demonstrated that senescence of AT2 cells (not loss of AT2) is sufficient to drive progressive pulmonary fibrosis. Others agree, and this is now an established pathophysiologic trigger in lung fibrosis(64, 124–126). Our findings are in keeping with published work (127) that suggests that AT2 senescence may be a targetable disease driver of lung injury in COVID-19. Although AT2 senescence is a shared phenomenon, our PPI network analyses --which integrated AT2 processes with the immune responses (ViP signatures) -- provided valuable clues into how platelet activation and thromboinflammation may be uniquely seen in the setting of COVID-19. These findings are in keeping with recent publications (128–130) which suggest that SARS-CoV-2 could induce epithelial senescence; senescent AT2 cells would then assume a SASP phenotype, which in turn led to neutrophil and platelet activation, and activation of the clotting cascade (128).

As for the limitations of this study, a direct comparison of lung samples from patients who survived COVID-19 but went on to develop (or not) restrictive (fibrotic) lung disease was not possible due to the lack of such datasets at this stage of the pandemic. It also remains unclear how to model post-COVID-19 progressive lung fibrosis *in vitro* and hence, no attempt was made to do so. Our results suggest that AT2 cells alone may be insufficient for such modeling because a specific host immune response that is carried in the PBMCs is a clear determinant as to who progresses and who does not. Although AT2-specific modulation of ER-stress pathway and SARS-CoV-challenged (treated vs untreated) hamsters were used to go beyond association and establish causation, our study did not attempt to inhibit/reverse fibrosis in COVID-19 by acting on any profibrogenic cellular pathway/process. Development of novel chemical matter/biologicals and validation of the therapeutic efficacy of such agents will take time, but if successful, our findings show that their benefits will likely transcend beyond PCLD into IPF and other fibrotic lung conditions such as IPF.

In conclusion, this transdisciplinary work provides insights into the pathogenesis of PCLD, formally defines the fibrogenic processes in the lung, and rigorously validates high value gene signatures or even targets (i.e., IL15, senescence pathways, etc.) to track and manipulate the same. The insights, tools, computationally vetted disease models, and biomarkers (prognostic gene signatures) identified here are likely to spur the development of therapies for patients with fibrotic interstitial lung disease of diverse causes, including IPF, all of whom have limited or no good treatment options.

## CONTRIBUTORS

DS, SS and PG conceptualized the project; SS and DS carried out all computational modeling and analyses and DS contributed all software used in this work; GDK carried out the human serum cytokine analysis; VC and SS carried out the immunohistochemical studies and their quantitative analysis under the supervision of PG; AGF carried out the immunofluorescence studies on COVID-19 lung tissues; SS carried out the PPI network creation and analysis; JD provided access to human subjects for cytokine analyses. CT, CRE and SD were responsible for the COVID-19 lung models. SS, DS and PG prepared figures for data visualization; SS and PG wrote the original draft of the manuscript; DS, SD, GK edited and revised the manuscript. DS, SD, and PG supervised various parts of the project and secured funding; DS and PG administered the project. SS, DS, and PG have accessed and verified the underlying data. All co-authors have approved the final version of the manuscript for submission.

## DATA SHARING STATEMENT

### Data availability

Source data are provided with this paper. All data is available in the main text or the supplementary materials. Publicly available datasets used are enlisted in **Supplemental Information 1**.

### Code availability

The software codes are publicly available at the following links: https://github.com/sahoo00/BoNE(131) and https://github.com/sahoo00/Hegemon (132). The PPI network analysis can be found at: https://github.com/sinha7290/PPIN.

## ACKNOWLEDGMENTS

This work was supported by the National Institutes for Health (NIH) grants R00-CA151673 and R01-GM138385 (to DS) and R01-AI141630, CA100768 and CA160911 (to P.G), R01DK107585 (SD) and R01-AI155696 (to P.G, D.S and S.D), UCOP-RGPO (R00RG2628 & R00RG2642 to P.G, D.S and S.D). GDK was supported through the American Association of Immunologists Intersect Fellowship Program for Computational Scientists and Immunologists. JMD was supported by U19AI142742. We thank Zea Borok (UCSD) for helpful discussions during the preparation of this manuscript.

