## Supplementary Materials for "COVID-19 lung disease shares driver AT2 cytopathic features with Idiopathic pulmonary fibrosis"

### SUPPLEMENTARY FIGURES

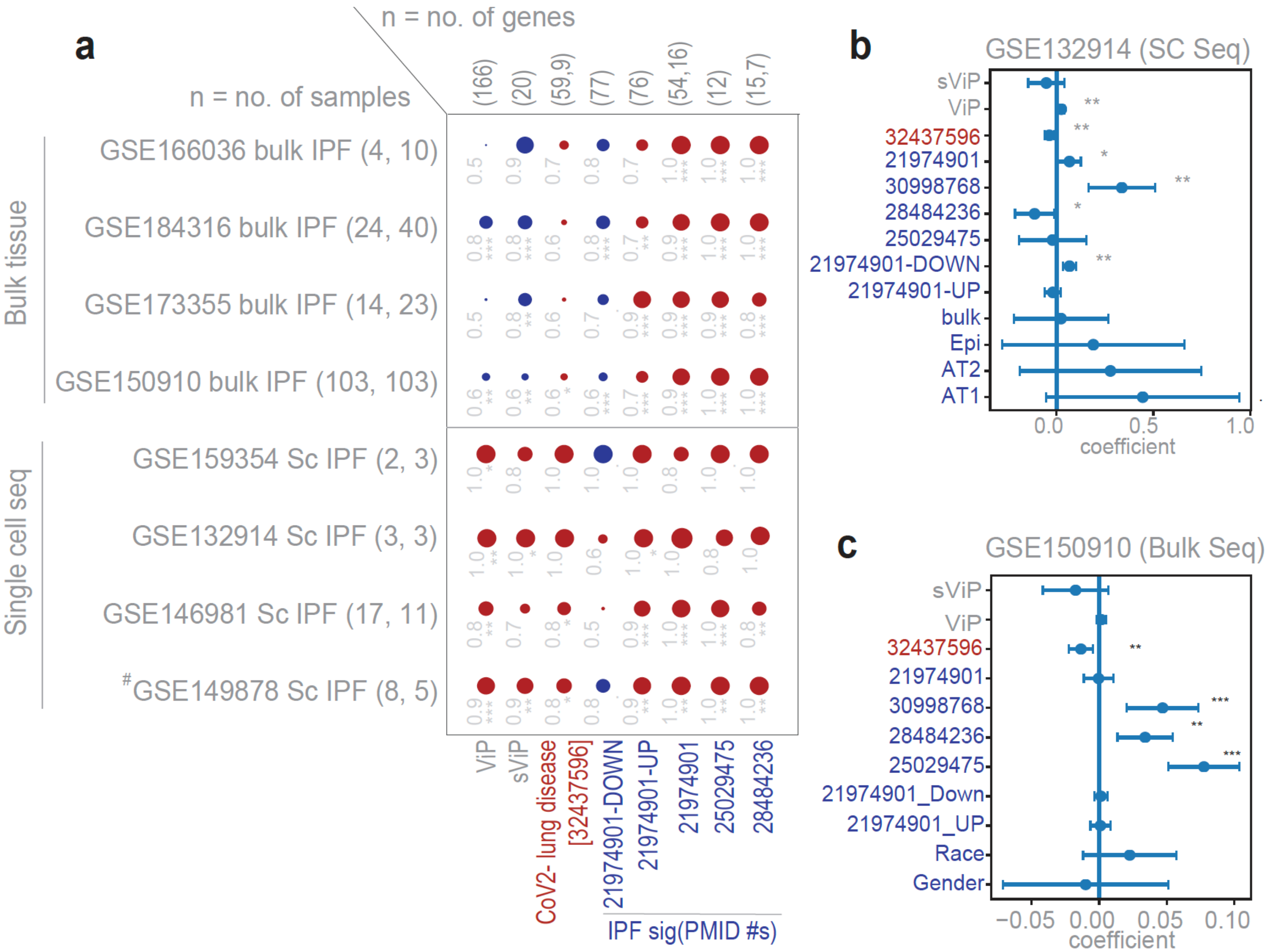


**Figure S1. Induction of both ViP and sViP signatures are detected in IPF by single cell RNA sequence (scRNA-seq) approach, but not by bulk tissue RNA sequence.**

**(a)** Bubble plots of ROC-AUC values (radii of circles are based on the ROC-AUC) demonstrating the direction of gene regulation (Up, red; Down, blue) in IPF lung-derived (i.e., lung explants) scRNA-seq datasets vs similar explant tissue-derived bulk RNA sequence datasets for ViP, sViP and other CoV and IPF signatures. GSE159354 and GSE149878 represent diverse cell types in the lung, whereas GSE132914 and GSE146981 are sorted epithelial cells only. ViP and sViP are prominently induced in single cell datasets but not in bulk tissue datasets. IPF signatures are generally induced to similar extent in all types of datasets. The significance of the AUC was determined by Welch's t-test ; * = $p < 0.05$, ** = $p < 0.01$, and *** = $p < 0.001$. (# denotes a pooled cohort of GSE149878; GSE122960).

**(b, c)** Multivariate analysis models the healthy vs IPF samples as a linear combination of composite scores of sViP and ViP (gray), CoV-lung (PMID# in red), and six independent IPF signatures (PMID#s in blue). For single cell dataset (**b**), cell type was also considered as a variable. The coefficient of each variable (at the center) with 95% confidence intervals (as error bars) and the *p* values were illustrated in the bar plot. The p-value for each term tests the null hypothesis that the coefficient is equal to zero (no effect). p-values are as follows: * = $p \leq 0.05$, ** = $p \leq 0.01$, and *** = $p \leq 0.001$.

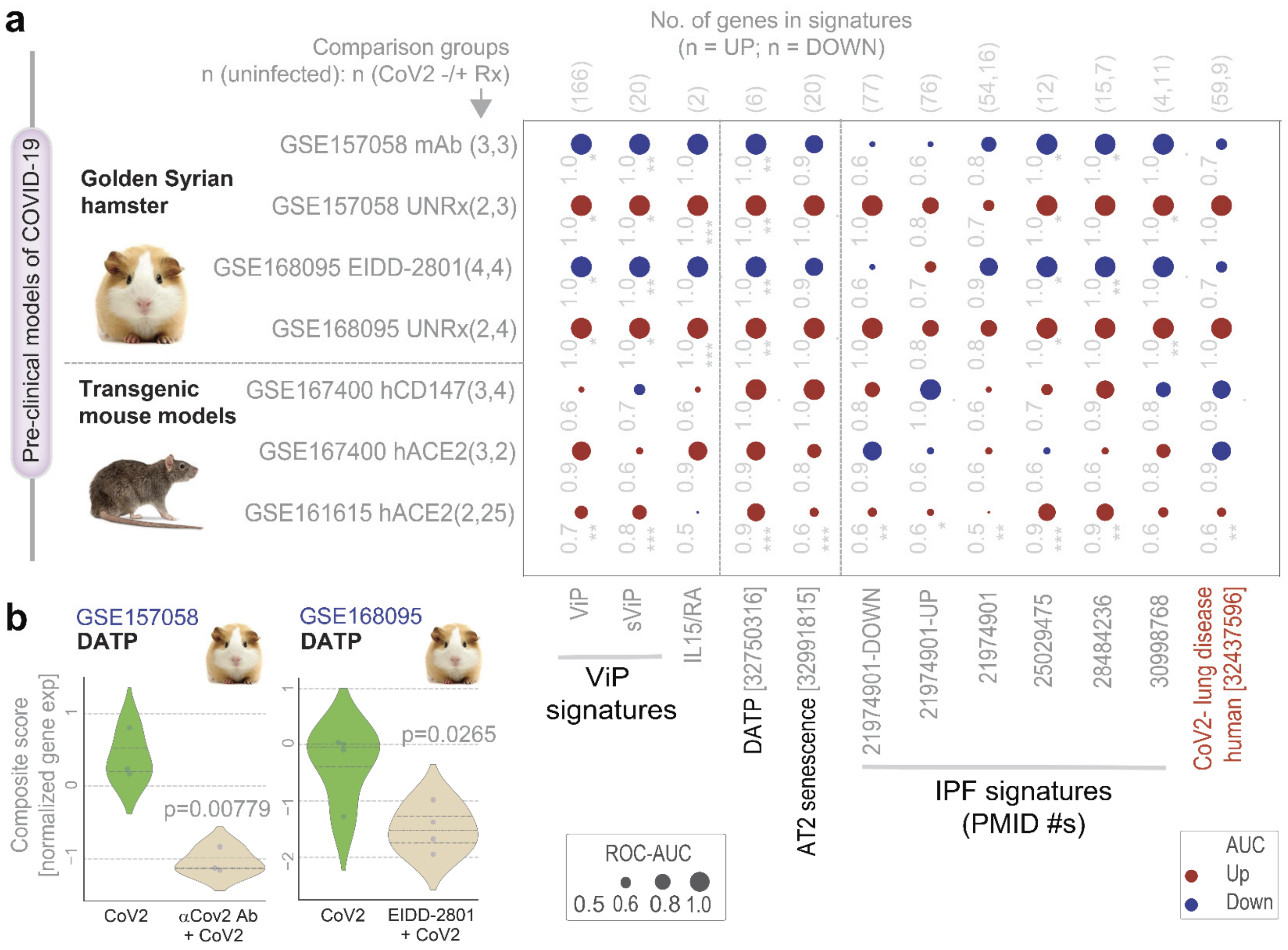


**Figure S2. Hamsters, but not humanized *tg*-mice faithfully recapitulate the host immune response and AT2 cytopathies that are found in COVID-19 and IPF.**

**a.** Schematic on the left summarizes the COVID-19 pre-clinical animal models analyzed in the bubble plots on the right. Bubble plots of ROC-AUC values (radii of circles are based on the ROC-AUC) demonstrating the direction of gene regulation (Up, red; Down, blue) for the classification of samples as follows (*from bottom to top*): uninfected vs. infected mouse lung samples (3 bottom rows), or uninfected vs. infected untreated (UNRx) hamster, or infected untreated vs. treated (with EIDD-2801(1) or Anti-Spike mAb(1)) hamster lung samples (4 top rows). The various signatures used for sample classification are listed below. **b**. Violin plots display the extent of induction of a KRT8+ AT2 damage-associated transient progenitor (DATP) signature(2). Welch's two sample unpaired t-test is performed on the composite gene signature score (z-score of normalized tpm count) to compute the *p* values. The significance of the AUC was determined by Welch's t-test ; * = p < 0.05, ** = p < 0.01, and *** = p < 0.001.

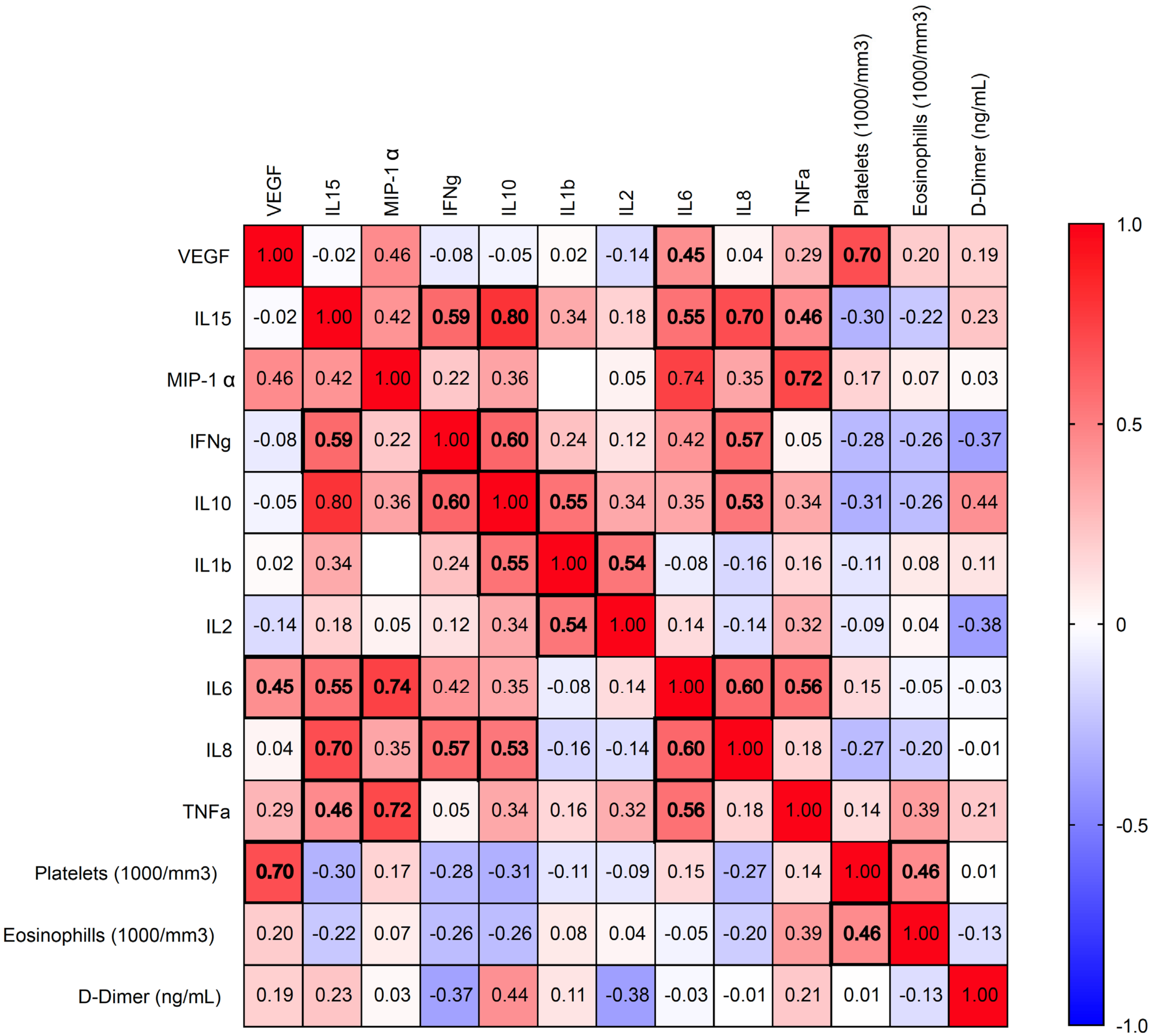


**Figure S3. Correlation matrix of cytokines levels in critical patients.** Pearson correlation tests of cytokine levels, as determined by mesoscale and clinical/laboratory findings. Heatmaps of correlation matrix are displayed for COVID19 critical patients (for patient information, see **Supplemental Information 3**). Visualization of data as matrix and determination of statistical significance were carried out using GraphPad Prism 9. Significant correlations, defined as those in which p values were < 0.05) are highlighted in bold fonts. Source data are provided as a **Supplemental Information 4.**
